# The Effect of Uncertainty on Prediction Error in the Action-Perception Loop

**DOI:** 10.1101/2020.06.22.166108

**Authors:** Kelsey Perrykkad, Rebecca P. Lawson, Sharna Jamadar, Jakob Hohwy

## Abstract

Among all their sensations, agents need to distinguish between those caused by themselves and those caused by external causes. The ability to infer agency is particularly challenging under conditions of uncertainty. Within the predictive processing framework, this should happen through active control of prediction error that closes the action-perception loop. Here we use a novel, temporally-sensitive, behavioural proxy for prediction error to show that it is minimised most quickly when variability is low, but also when volatility is high. Further, when human participants report agency, they show steeper prediction error minimisation. We demonstrate broad effects of uncertainty on accuracy of agency judgements, movement, policy selection, and hypothesis switching. Measuring autism traits, we find differences in policy selection, sensitivity to uncertainty and hypothesis switching despite no difference in overall accuracy.

---

A significant challenge to an agent’s perceptual and decision-making processes is to distinguish between sensations that it can control, and those out of its control. For example, imagine you are working on your computer and it beeps. How do you know if you caused it, as opposed to a colleague emailing you? Influential theoretical work on predictive processing and active inference suggests that the brain relies on prediction errors to assess and test hypotheses about agency (Friston et al., 2013), but empirical evidence for this suggestion is lacking.

Inferring the relations between actions and their sensory consequences is riddled with uncertainty due to the complexities involved in deconstructing sensory evidence from the non-linear confluence of hidden causes. Sometimes when you click, the ensuing beep occurs later because the computer is updating its virus-software; other times, it happens straight away. The brain must represent this uncertainty at numerous hierarchical levels to identify when it is appropriate to attribute agency to oneself. In this example, the breadth of the distribution representing how long it takes for the beep to occur is the *variability* and the frequency of the virus-updates is the *volatility* (how often does the variability distribution change). Crucially, we do not yet know how this uncertainty changes ongoing decisions about *which* actions to perform when trying to explore and infer agency; thus, we have yet to explore how agents close the action-perception loop under uncertainty.

A *judgement of agency* is the verdict that the agent was herself the source of a sensory event – the conscious “I did that” response. It is often (but not always) based on a *sense of agency* (or a feeling of authorship) during the movement. Predictability is often investigated in sense and judgement of agency paradigms by manipulating whether or not the identity (Bednark, Poonian, Palghat, McFadyen, & Cunnington, 2015; Engbert & Wohlschlager, 2007; Hughes, Desantis, & Waszak, 2013; Kuhn et al., 2011; Majchrowicz & Wierzchoń, 2018), timing (Hughes et al., 2013; Majchrowicz & Wierzchoń, 2018) and/or presence (Moore & Haggard, 2008) of a sensory outcome meets some prediction set up by the block-wise probability of each outcome. However, very few studies consider a more continuous distribution of deviations from the expected outcome (e.g., Zalla, Miele, Leboyer, and Metcalfe (2015)) and, to our knowledge, no previous studies have considered volatility (changes to such a distribution) in an agency paradigm.

In classic agency experiments, there are so few actions available to participants that action-selection strategies (or *policies*) cannot easily change in response to changes in prediction error or uncertainty. In some designs, such as Desantis, Hughes, and Waszak (2012), specific actions trigger specific outcomes, but the participants are instructed to equally perform each action. This does not allow participants to explore and attempt to optimally vary policy-selection. In other studies, participants do have freedom to change strategy, and have online action outcome mismatches, but the dependent variables are not sensitive to these strategies and so the temporal dynamics of online decisions with respect to this error are unknown (Zama, Takahashi, & Shimada, 2017). This gap in knowledge is crucial for understanding how we distinguish self-generated and externally-caused sensations in the real world. The current study sought to close this gap using a novel judgement of agency task that dynamically closed the action-perception loop while independently manipulating variability and volatility.

In the current study, forty human participants made freeform mouse movements to identify which (if any) of eight moving squares they controlled. Each block of trials had high or low variability and high or low volatility (both within-trial). For a schematic diagram of the experimental set up, task and experimental manipulation, see **Figure 1** and a video of the task is available at https://figshare.com/s/fd2742b897e21d901dd0 (DOI: 10.26180/5eabbfb9a8aa4). Full details in *Methods* below.

**Figure 1.**
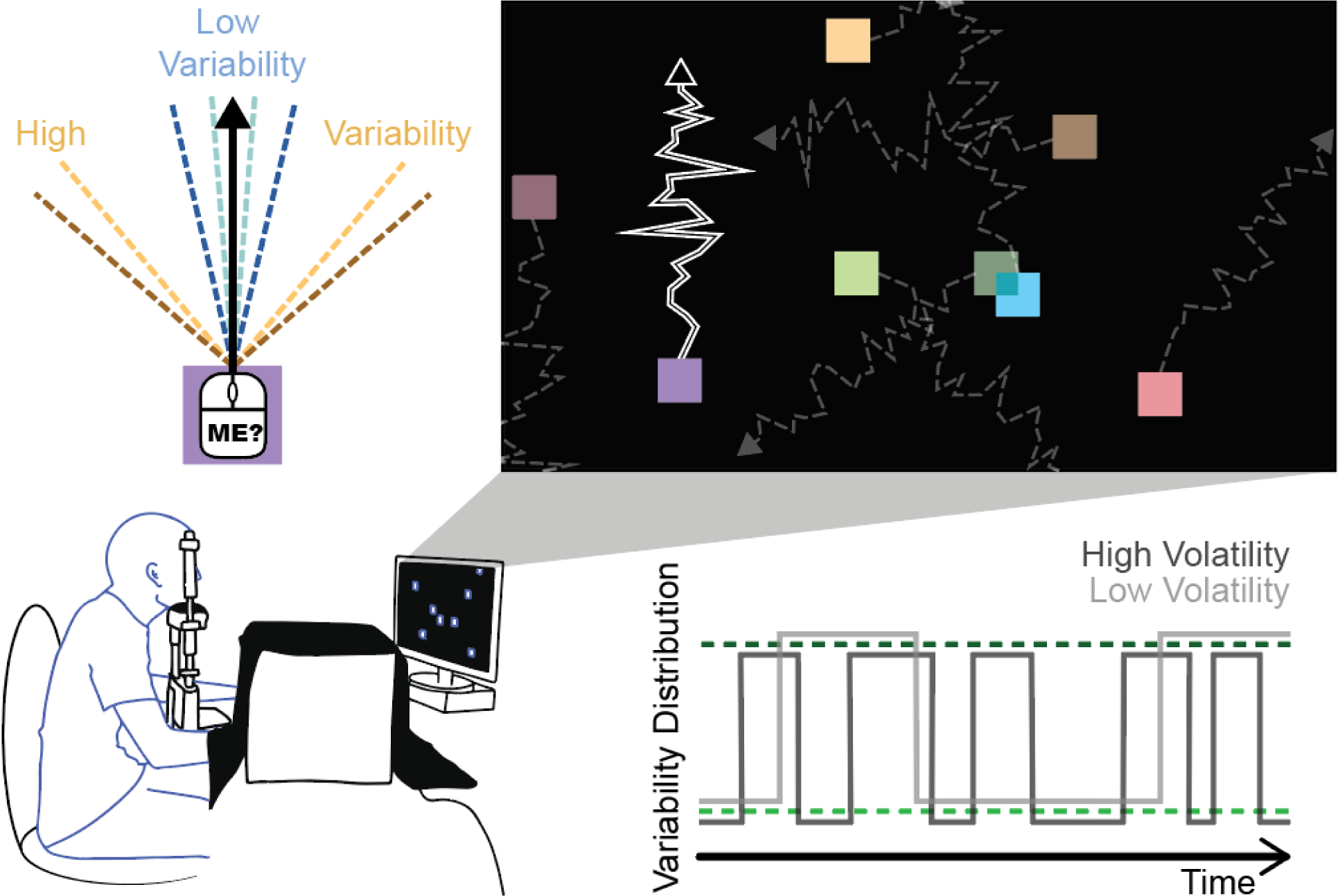
Task and Manipulation. The participants’ task involved using a hidden mouse to control eight squares on the screen. The mean of the target square’s movement was the participants’ movement, and distractor squares moved at a random angle offset from mouse movement. Jitter was added to the motion of all squares depending on the condition. In *low variability* blocks, the 95% confidence interval from which jitter was sampled switched between 10° (light blue) and 30° (dark blue), and for *high variability*, it switched between 90° (yellow) and 110° (brown) around the input movement or offset. In *low volatility* the distribution changed three random times per trial (light grey), and in *high volatility* it changed ten times (dark grey). See also https://figshare.com/s/fd2742b897e21d901dd0 (DOI: 10.26180/5eabbfb9a8aa4) for a video demonstration.

To understand these missing components in the process of inferring agency, we turn to recent accounts of agency from predictive processing - an explanatory framework whose fundamental claim is that the brain’s function is to minimise the long-term average error between its expected and actual sensory input (*prediction error*) (Clark, 2015; Friston, 2010; Hohwy, 2013). In doing so it reduces uncertainty by refining models of the hidden causes of sensory input in the environment and in the agent itself.

Prediction error can be minimised by updating expectations while passively receiving sensory input (*perceptual inference; perception*). Another way to minimise prediction error is through action, by selectively sampling sensory input to satisfy beliefs about sensory input in future states of the world and the agent’s own body, given certain actions (*active inference; action*) (Friston, 2017). Previous agency research has focused on perceptual inference in the context of agency, and has not interrogated the ongoing process of active inference.

Under an active inference account, agency attribution would occur by minimising the divergence between the predicted outcomes of available policies for action and the most probable future sensory states; in other words, when there is a belief that goals can be reached from the agent’s current state (Friston, Samothrakis, & Montague, 2012; Friston et al., 2013; Hohwy, 2015). Thus, precision (i.e., the inverse of uncertainty) of these inferences is important (Friston et al., 2013) and lead us to investigate the effect of such variability on actions, prediction error and inferred agency.

According to active inference, the very purpose of action is then to minimise expected prediction error. To understand how this plays out in the action-perception loop it is then essential to reveal the interplay between action selection and the magnitude of prediction error at a given time, under a given policy. For the critical case of agency attribution, it is not known how an agent infers policies that may help reduce uncertainty about agency; this is mainly because thus far its magnitude has been under the control of experimenters, not participants themselves. Here, rather than dictating the magnitude of prediction error and measuring effects on behaviour and neural processes, we instead measure the prediction error itself and allow participants to control it with their actions.

The most straightforward expectation for active interrogation in an action-perception loop is that, where possible, policies are inferred which minimise prediction error. Part of the difficulty in testing this prediction is finding an appropriate way to measure prediction error. Here, we operationalise prediction error using eye position to calculate the evolving divergence between hand-movement and stimulus trajectories. Eye-tracking indicates moment-to-moment beliefs about agency which can be tested by mouse-movement. We predict that variability and volatility will have independent effects on movement patterns and policy selection, as well as on prediction error minimisation and subsequent judgements of agency. Specifically, high variability allows less precise representation of control states, which predicts more repetitive policy selections (Perrykkad & Hohwy, 2020), more prediction error and less accurate judgements of agency. High volatility suggests potentially discoverable interfering hidden causes, predicting more policy exploration and more variance in prediction error which could aid accurate inference of agency. Independent of accuracy, we expect a positive correlation between agency-driven prediction error minimisation and judgements of agency, partly based on active inference theory and partly on prior literature on the role of prediction-expectation mismatch for agency reports.

It is instructive to consider how prediction error minimisation might differ in clinical or subclinical populations because such comparisons help reveal how the prediction error mechanisms work. We focus here on predictive processing accounts of autism, according to which autistic individuals have difficulty abstracting causal rules to higher statistical levels, and thus classify more uncertainty as irreducible. This has been theorised to be due to weaker priors, weightier prediction errors or hyper-flexible estimates of volatility, which all result in a higher learning rate in autism (for review and details, see (Palmer, Lawson, & Hohwy, 2017)). Manipulation of uncertainty in tasks that rely on perceptual inference has been shown to change performance in autistic populations (Lawson, Mathys, & Rees, 2017). Characteristic differences in action in autism, such as restricted and repetitive behaviours, may indicate differences in active inference in variable environments (Palmer et al., 2017). Previous research, not framed in terms of predictive processing, have used basic versions of the task we use here, and found no difference between groups of autistic and non-autistic participants (Grainger, Williams, & Lind, 2014; Russell & Hill, 2001; Williams & Happé, 2009), however, we predict the relationship between autism traits and agency attribution should be specific to interactions with uncertainty in the environment as the action-perception loop is dynamically closed (cf. Zalla et al. (2015)). This in turn speaks to underexplored topics in autism research relating to the sense of self and agency (Perrykkad & Hohwy, 2019). Hence, here we additionally measured autistic traits in our sample and we predict that uncertainty will differently affect policies for movement and prediction error minimisation for participants along this scale.

## Results

In this section, we summarise all statistical models in three sections, first, covering prediction error measures, second, the movement and strategy measures and last, overall task performance. For each section, we describe the effect of uncertainty on the dependent variables followed by AQ results (though all statistical models included all fixed factors as above). For brevity, we report only main effects of uncertainty in the movement and policy variables. Full statistical reporting is included in Supplementary Materials. See also Table 1 for a summary of all significant results.

**Table 1.**
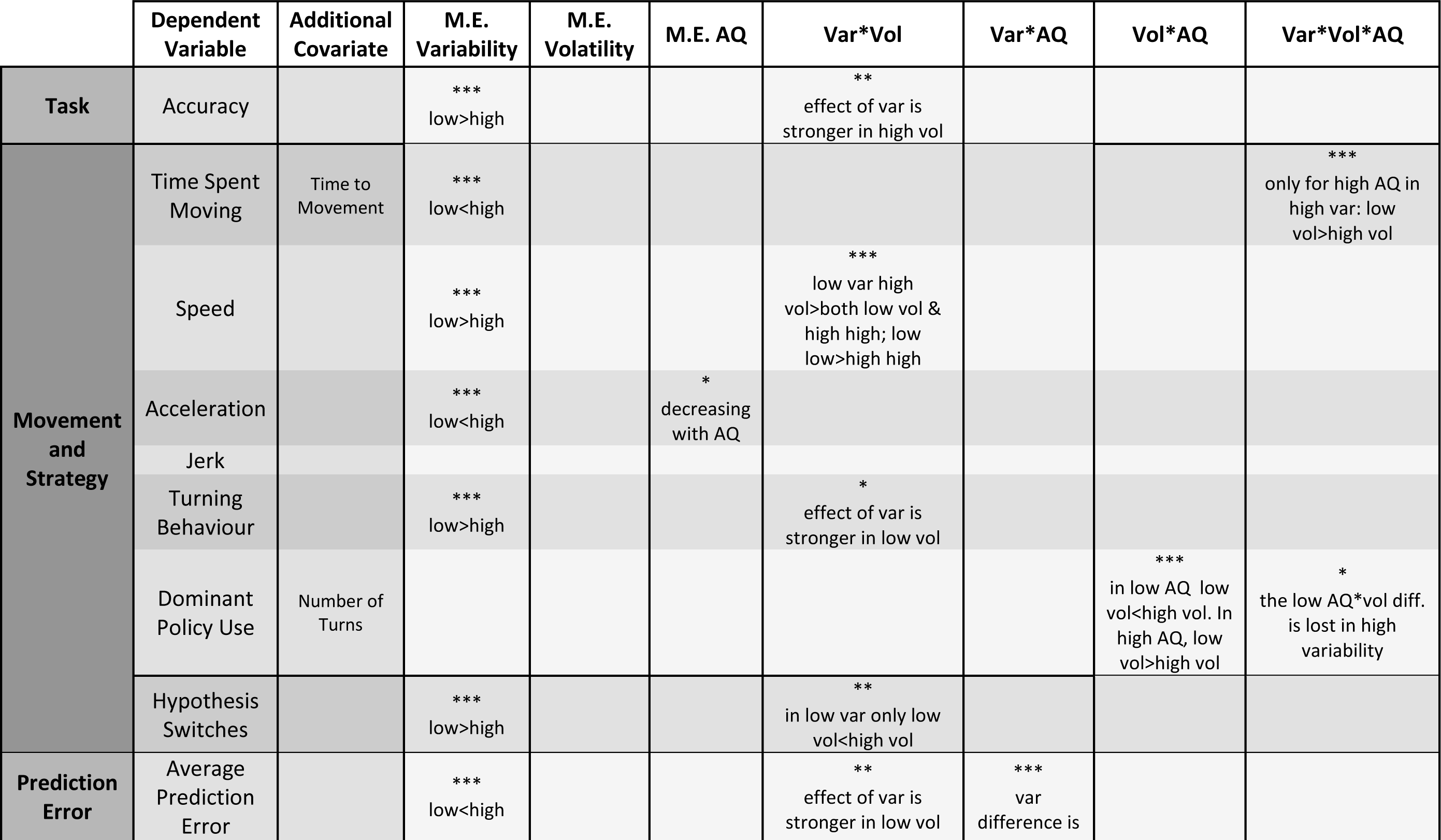
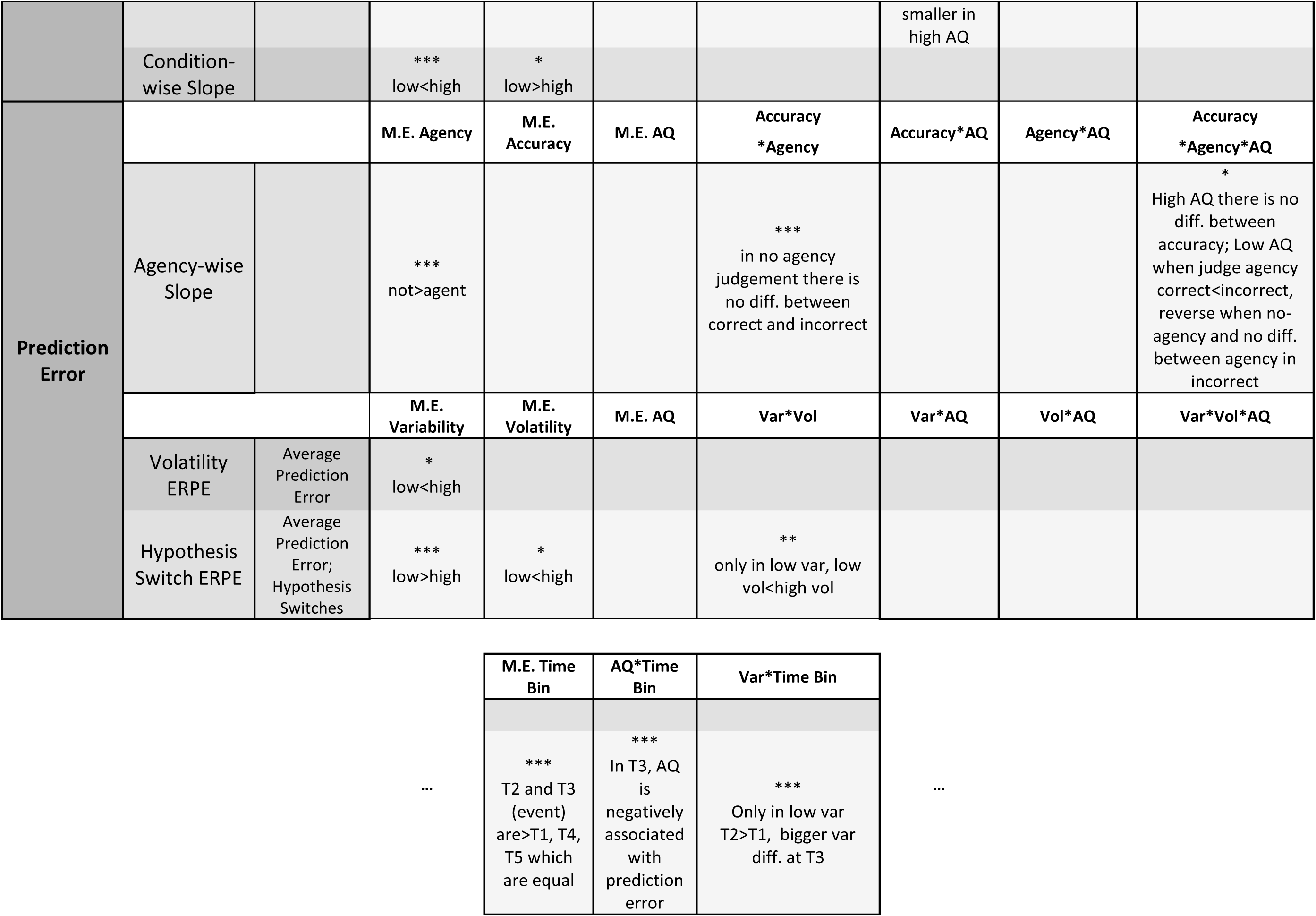
Significant Results Summary: For variables, M.E.=Main Effect, *=Interaction, For results, *=p ≤ 0.05, **=p ≤ 0.01, ***=p ≤ 0.001 (Post-hoc values Bonferroni corrected for multiple comparisons), Var=Variability, Vol=Volatility, AQ=Autism Quotient, T1-5=Time bins 1-5, diff.=difference

|  | Dependent Variable | Additional Covariate | M.E. Variability | M.E. Volatility | M.E. AQ | Var*Vol | Var*AQ | Vol*AQ | Var*Vol*AQ |
| --- | --- | --- | --- | --- | --- | --- | --- | --- | --- |
| **Task** | Accuracy |  | ***<br>low>high |  |  | **<br>effect of var is stronger in high vol |  |  |  |
| **Movement and Strategy** | Time Spent Moving | Time to Movement | ***<br>low<high |  |  |  |  |  | ***<br>only for high AQ in high var: low vol>high vol |
|  | Speed |  | ***<br>low>high |  |  | ***<br>low var high vol>both low vol & high high; low low>high high |  |  |  |
|  | Acceleration |  | ***<br>low<high |  | *<br>decreasing with AQ |  |  |  |  |
|  | Jerk |  |  |  |  |  |  |  |  |
|  | Turning Behaviour |  | ***<br>low>high |  |  | *<br>effect of var is stronger in low vol |  |  |  |
|  | Dominant Policy Use | Number of Turns |  |  |  |  |  | ***<br>in low AQ low vol<high vol. In high AQ, low vol>high vol | *<br>the low AQ*vol diff. is lost in high variability |
|  | Hypothesis Switches |  | ***<br>low>high |  |  | **<br>in low var only low vol<high vol |  |  |  |
| **Prediction Error** | Average Prediction Error |  | ***<br>low<high |  |  | **<br>effect of var is stronger in low vol | ***<br>var difference is smaller in high AQ |  |  |
|  | Condition-wise Slope |  | ***<br>low<high | *<br>low>high |  |  |  |  |  |
| Prediction Error |  |  | M.E. Agency | M.E. Accuracy | M.E. AQ | Accuracy<br>*Agency | Accuracy*AQ | Agency*AQ | Accuracy<br>*Agency*AQ |
|  | Agency-wise Slope |  | ***<br>not>agent |  |  | ***<br>in no agency judgement there is no diff. between correct and incorrect |  |  | *<br>High AQ there is no diff. between accuracy; Low AQ when judge agency correct<incorrect, reverse when no-agency and no diff. between agency in incorrect |
|  |  |  | M.E. Variability | M.E. Volatility | M.E. AQ | Var*Vol | Var*AQ | Vol*AQ | Var*Vol*AQ |
|  | Volatility ERPE | Average Prediction Error | *<br>low<high |  |  |  |  |  |  |
|  | Hypothesis Switch ERPE | Average Prediction Error; Hypothesis Switches | ***<br>low>high | *<br>low<high |  | **<br>only in low var, low vol<high vol |  |  |  |

|  |  |  |  |  |
| --- | --- | --- | --- | --- |
|  | M.E. Time Bin | AQ*Time Bin | Var*Time Bin |  |
| ... | ***<br>T2 and T3 (event) are>T1, T4, T5 which are equal | ***<br>In T3, AQ is negatively associated with prediction error | ***<br>Only in low var T2>T1, bigger var diff. at T3 | ... |

### Prediction Error

Across all participants, the average calculated prediction error per trial was 10.5 pixels per frame (**Figure 2a;** σ=4.93, Range=0.43-50.1). An MLM analysis revealed that, as expected, average prediction error across each trial was significantly associated with the variability condition (F(1,4661)=742.63, p<0.001). Additionally, there was a significant interaction between variability and volatility (F(1,4661)=5.14, p=0.023). Post-hoc analyses showed that this was because the effect of variability was stronger under low volatility than high (z>17.80, p<0.001).

**Figure 2.**
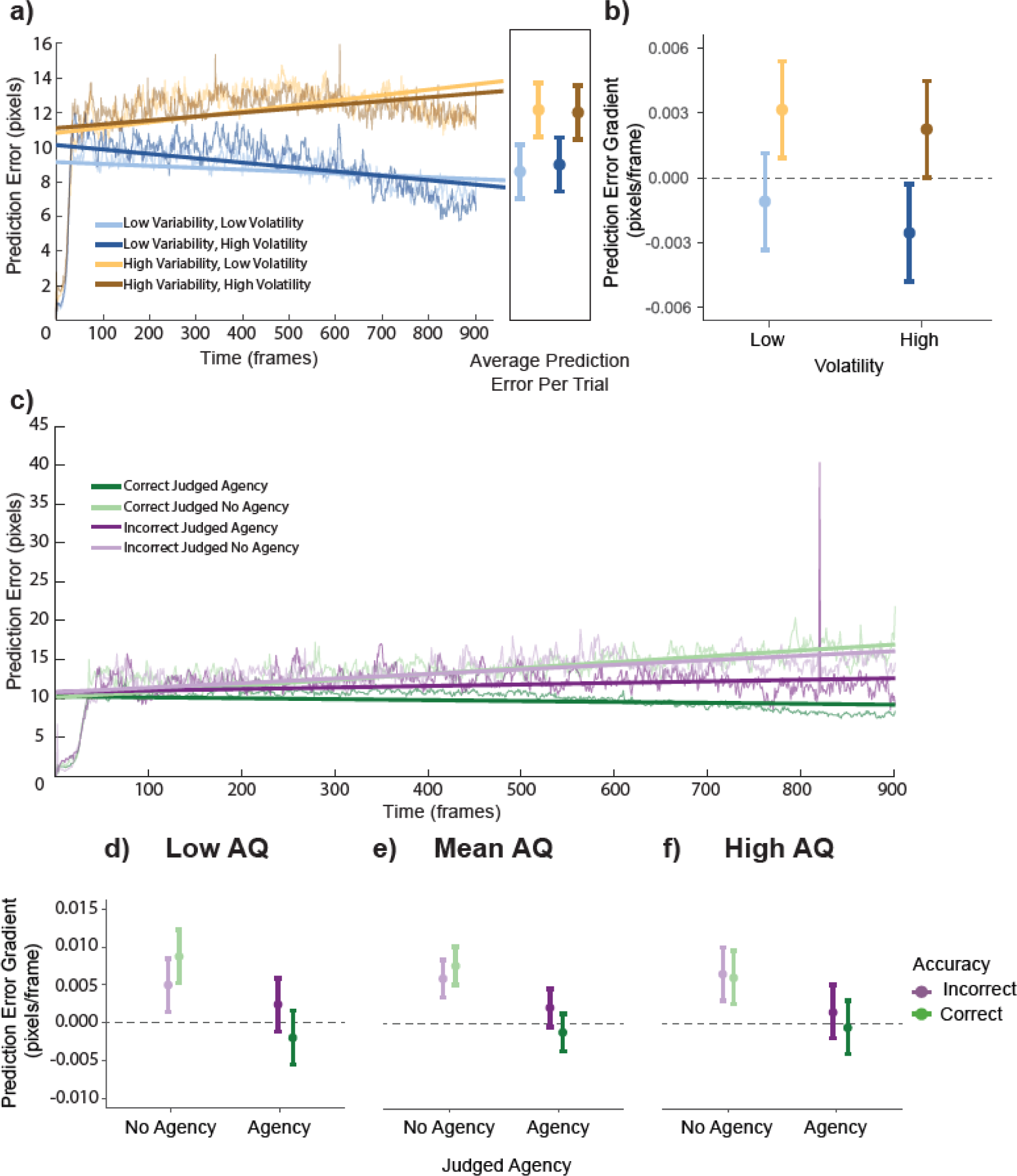
Prediction Error Average and Gradient. Panel a) shows the grand average prediction error across the trial split by condition with lines of best fit for each. The box at the end of the graph shows the average prediction error across trials in each condition. Panel b) shows the mean gradient or slope for the lines of best fit for each participant in each condition. Panel c) shows the grand average prediction error across the trial split by correct (green) and incorrect (purple) trials and whether the participants chose a square (Judged Agency, dark colours) or said that it was a no-control trial (Judged No Agency, light colours) with lines of best fit for each. Panels d-f show the mean gradient or slope for the lines of best fit for each participant in each combination, split by AQ score. Error bars are 95% CI.

Comparing the slope of prediction error in each condition, an MLM revealed a significant main effect of variability (F(1,114)=58.15, p<0.001), which indicated that there was more prediction error minimisation in the low variability condition (lower gradient) than the high (See **Figure 2a and 2b**; t=7.63, p<0.001). There was also a marginally significant main effect of volatility (F=3.96, p=0.049), which showed a trend toward more prediction error minimisation in high volatility compared to low volatility (t=1.99, p=0.049).

To investigate the relationship between prediction error minimisation and the participant’s judgements, we performed an MLM with a different structure. For each participant, a linear fit to prediction error across trials with the same accuracy and agency judgement served as the dependent variable. AQ score, accuracy and agency were included as fixed effects, and participant as a random intercept. This MLM showed a main effect of agency (F(1,113)=82.89, p<0.001) and an interaction between agency and accuracy (**Figure 2c**; F(1,113)=12.79, p<0.001). Agency was associated with increased prediction error minimisation (t(113)=9.10, p<0.001). Post-hoc tests for the interaction showed that only when participants judge that they did not control any of the stimuli was there no difference in prediction error minimisation between correct and incorrect trials (t(113)=1.74, p=0.51). When participants judge that they did have agency, there is more prediction error minimisation when they are correct than incorrect (t(113)=3.31, p=0.007). Numerically, the mean slope of the prediction error was only negative (indicating successful prediction error minimisation) when participants were both accurate and judged that they had agency.

To look at the effect of uncertainty and AQ score on dynamics of prediction error and hypothesis testing, we performed an MLM on the ERPE centered on hypothesis switches. In addition to the standard MLM, we included time-bin as an additional fixed effect of interest and average prediction error and average number of hypothesis switches in each condition as fixed-effect covariates. **Figure 3a** shows the timeseries for the average prediction error across conditions and participants in the analysed epoch. There were significant main effects of variability (F(1,512)=125.10, p<0.001), volatility (F(1,719)=6.14, p=0.013) and time-bin (F(4,719)=252.29, p<0.001) and two-way interactions between variability and volatility (**Figure 3b**; F(1,729)=10.76, p=0.001) and variability and time-bin (F(4,719)=17.94, p<0.001). Time bins one, four and five were not significantly different from one another (t(720)=0.06-1.55, p=1.00) but that the others were all significantly different from one another (t(720)=5.24-26.79, p<0.001), indicating a significant increase before the hypothesis switch starting at least 300ms before, and a drop after back to the initial level of prediction error. Post-hoc analyses into the main effect of variability showed that low variability conditions had *greater* prediction error around the time of a hypothesis switch than did high variability conditions (t(519)=11.09, p<0.001), which is the inverse of the pattern for average prediction error across the whole trial. Post-hoc analysis of the interaction between time-bin and variability showed that the difference between variability conditions held across all time bins surrounding the hypothesis switch (t(697)=6.15—13.81, p<0.001), but that this difference was greater during time bin three (3.89 pixels, greater than other bin averages by at least 1.69 pixels). Further, only in low variability is there a significant increase from time bin one to two (t(720)=6.15, p<0.001), indicating the increase may occur closer to the event in high variability conditions. While overall, low volatility was associated with less prediction error than high volatility around the time of hypothesis switches (t(721)=2.48, p=0.013), post-hoc analysis of the interaction between variability and volatility showed that this only holds when variability was low (t(727)=4.07, p<0.001).

These findings suggest that increased prediction error minimisation is associated with reduced environmental variability but increased volatility, and correctly and positively inferring agency. We have also shown that hypothesis switches function to reduce rising prediction error, and that the dynamics of minimising prediction error in this way is affected by environmental uncertainty.

**Figure 3.**
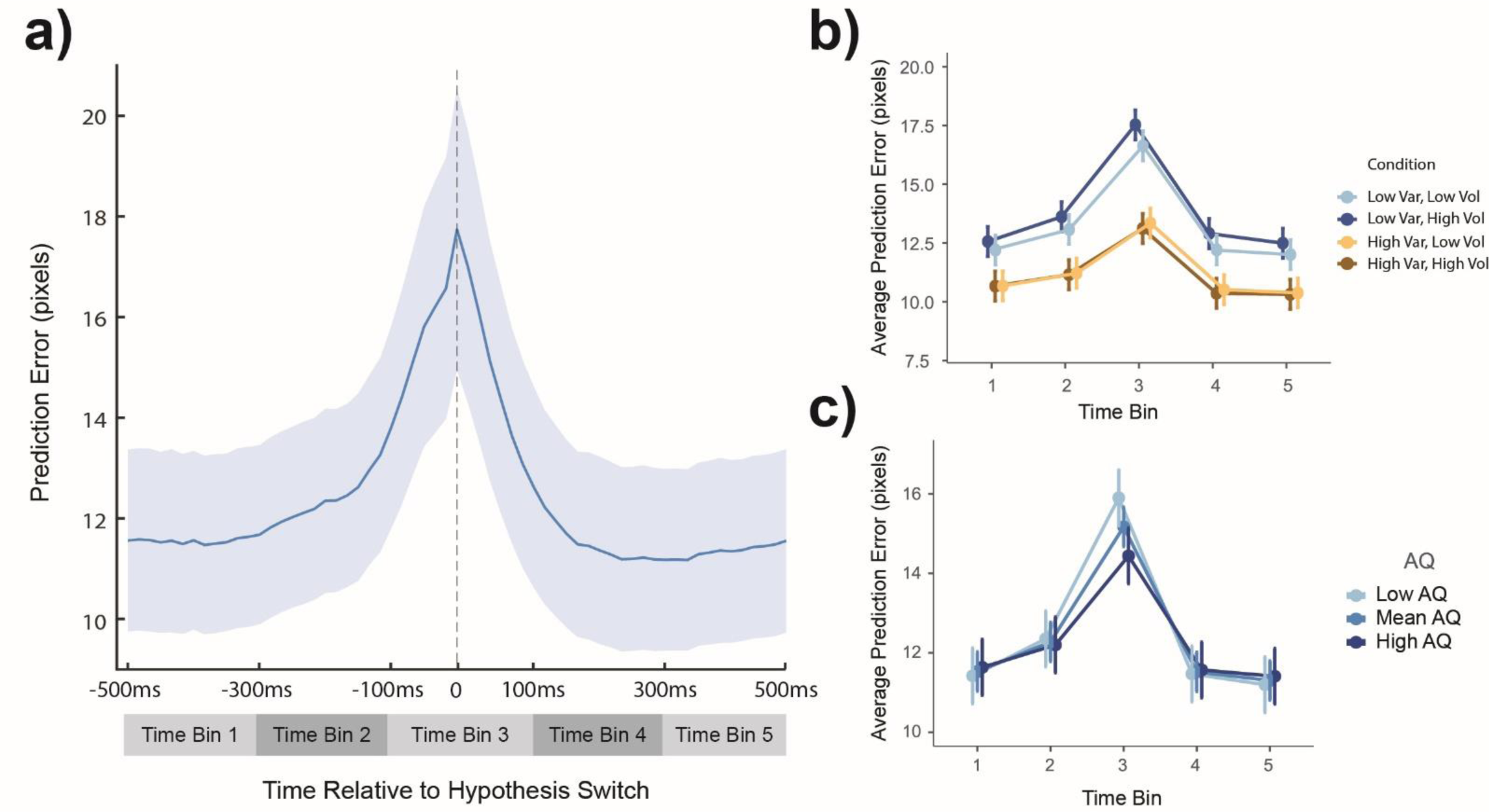
Hypothesis Switch Event-Related Prediction Error (ERPE) Panel a) shows the grand average (blue line) prediction error across participants in a one second epoch centered on hypothesis switches. Time bins used for statistical models are represented in grey shaded bars below. Data used in statistical models, and therefore in panels b) and c) is adjusted for average prediction error differences between conditions and average number of hypothesis switches. Panel b) shows average prediction error in each time bin for each condition. There is more prediction error in this epoch in low variability (blue) conditions than high (orange). This difference is greatest in time bin three, at the time of the event. In low variability (blue), low volatility (light blue) conditions showed less prediction error in this epoch than high (dark blue). Time bin three has the greatest prediction error, followed by time bin two, and none of the others are significantly different from each other. The increase from time bin one to two is only significant in low variability (blue). Panel c) shows the data split by AQ score - lower AQ scores (lightest blue) are associated with greater prediction error at the time of the event (time bin three). Error bars and shading are 95% CI.

### Autism Traits and Prediction Error

The model considering the effect of uncertainty and autism traits on average prediction error across a trial showed a significant interaction between variability and AQ (F(1,4661)=31.74, p<0.001). Post-hoc analyses of the variability x AQ interaction showed that the difference between variability conditions decreases as AQ increases (though they are still significantly different across all AQ scores; z=15.28-27.25, p<0.001).

Additionally, the MLM considering agency, accuracy and AQ showed a three-way interaction between these variables (F(1,113)=5.69, p=0.02). Post-hoc analyses showed no difference between agency judgements for incorrect trials for participants with a low AQ score (**Figure 2d**; F(1,113)=3.51, p=0.064), but otherwise, when participants judged that they had agency over one of the stimuli, the slope of their prediction error was lower, indicating that they were more effective at minimising prediction error (t(113)=9.10, p<0.001) (both the mean and high AQ groups, and when correct in low AQ). Further, while low AQ participants’ prediction error was maximally sensitive to accuracy (lower slopes when correctly judging agency than incorrectly doing so, F(1,113)=10.06, p=0.002; and lower slopes when incorrectly denying agency than when correctly doing so F(1,113)=7.75,p=0.006); high AQ participants’ prediction error was not sensitive to accuracy at all (**Figure 2f**; F(1,113)=0.11-2.29, p=0.13-0.74). Participants with a mean AQ showed the appropriate difference only when they judged that they had agency (F(1,113)=10.99, p=0.001, F=10.99).

Looking at the prediction error dynamics limited to the epoch around hypothesis switches showed a significant interaction between AQ and time-bin (**Figure 3c**; F(4,719)=12.16, p<0.001). Post-hoc analysis showed a significant difference only in time-bin three (the time of the event) depending on the AQ score (F(1,50)=8.58, p=0.005). A further Pearson’s correlation test of AQ by prediction error in this time-bin showed that as AQ increased, the prediction error at the time of a hypothesis switch decreased (r=-0.21, p<0.001).

These findings suggest that uncertainty in the environment differentially affects participants’ prediction error depending on measured autism traits, including the relationship between prediction error minimisation and judgement of agency, and propensity to switch hypotheses in response to increasing prediction error.

### Movement Characteristics and Policy Selection

Participants moved for an average of 13.7 seconds per trial (σ=1.55, Range=3.95-15.1). An MLM comparing the average duration of each trial spent moving across conditions (with the additional fixed effect of time to movement on each trial to account for possible confound) found a significant main effect of variability (**Figure 4a;** F(1,4660)=727.71, p<0.001). Participants moved for longer in high variability conditions compared to low variability conditions by an average of 801ms.

**Figure 4.**
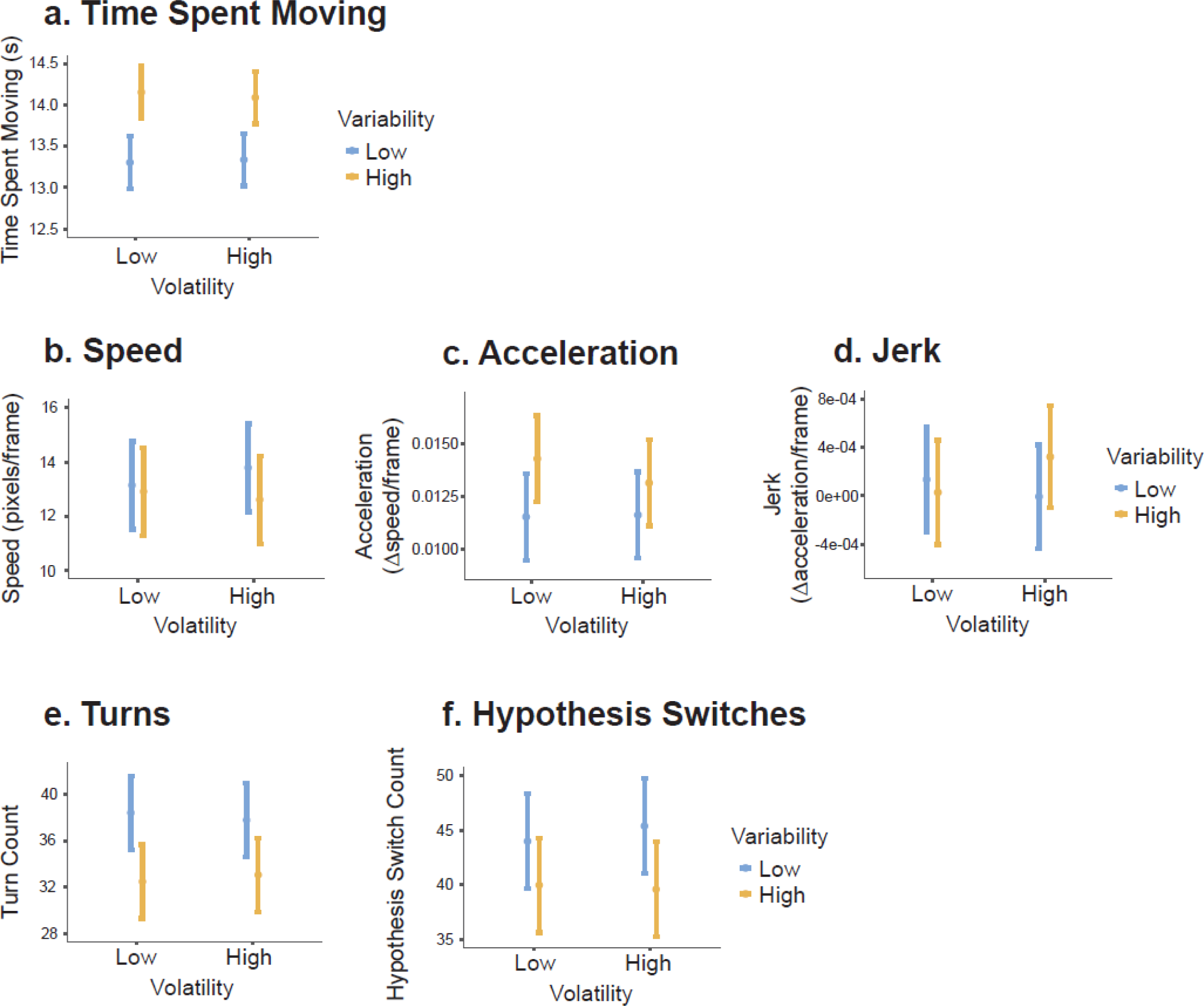
Movement and Strategy. These graphs depict movement and strategy variables (except dominant policy use, see **Figure 5**) across all participants. Volatility is along the x-axis for each graph. Orange bars represent high variability, blue bars represent low variability. Error bars are 95% CI. a) shows mean duration of each trial spent moving, controlling for time to movement onset on each trial. b) shows average speed of movement, c) average acceleration and d) average jerk. e) shows average turn count on each trial. f) shows the average number of hypothesis switches on each trial, when the participant moves their eyes from one square to another.

An MLM analysis on average speed of movement revealed a significant main effect of variability (**Figure 4b**; F(1,4661)=36.42, p<0.001) such that participants moved faster in the low variability condition compared to high (z=6.03, p<0.001). An MLM on acceleration showed a main effect of variability (**Figure 4c**; F(1,4664)=12.68, p<0.001), with faster average acceleration in the high variability trials, compared to low (z=3.56, p<0.001). An MLM on jerk showed no significant results (**Figure 4d**).

On average, each trial contained 35 turns (**Figure 4e**; σ=13.9, Range=6-107). An MLM on turn count showed a significant main effect of variability (F(1,4661)=346.22, p<0.001) such that participants turned more frequently in low variability than high variability trials (z=18.61, p<0.001). The dominant turn-types (*policies*) in order of frequency across participants were Non-Cardinal (n=22), Hesitant-straight (n=14), Horizontal (n=3) and Circle (n=1). On average, in each trial, participants used their dominant policy 39.3% of the time (σ=16.0, Range=0-100), and within each participant, the average percent of turns on each trial that were of their dominant policy ranged from 30.1% to 51.9%. For the MLM on dominant policy turn count for each trial, the additional covariate of absolute number of turns on each trial was included to account for this confound. There were no significant main effects.

On average participants switched hypotheses 42.2 times per trial (**Figure 4f**; σ=13.48, Range=6-134). An MLM on hypothesis switch counts in each trial showed a main effect of variability (F(1,4661)=195.91, p<0.001) such that there are more switches in low variability than in high (z=14.00, p<0.001).

These findings suggest that participants’ movement was strongly affected by increased environmental variability, causing participants to move more, move slower but accelerate more quickly, and switch hypotheses less often.

### Autism Traits and Movement and Policy

For the dependent variable of time spent moving, there was a significant three-way interaction between AQ, variability and volatility (F(1,4660)=11.37, p<0.001). Post-hoc tests showed that for participants with high AQ only, under high variability only, participants moved for an average of 200ms longer in low volatility than high volatility conditions. Additionally, the model considering acceleration showed a main effect of AQ (F(1,38)=5.73, p=0.022), such that mean acceleration decreased with AQ (R^2^=-0.011, p<0.001). There were no significant findings relating to autism traits across other movement characteristics.

Considering how participants across the AQ range changed their policies in response to uncertainty, the model for dominant policy use showed a significant interaction between AQ and volatility (F(1,4660)=19.17, p<0.001), and a significant three way interaction between AQ, variability and volatility (**Figure 5**; F(1,4661)=4.27, p=0.039). Post-hoc analyses for the two-way interaction showed that for low AQ (**Figure 5a**) participants used their dominant policy more in the high volatility condition (z=2.14, p=0.032), but only when variability was low (z=2.92, p=0.004), otherwise volatility made no difference (z=0.10, p=0.918). For high AQ (**Figure 5c**) participants used their dominant policy more in the low volatility condition (z=4.05, p<0.001), regardless of the variability (high: z=2.20, p=0.028; low: z=3.53, p<0.001).

**Figure 5.**
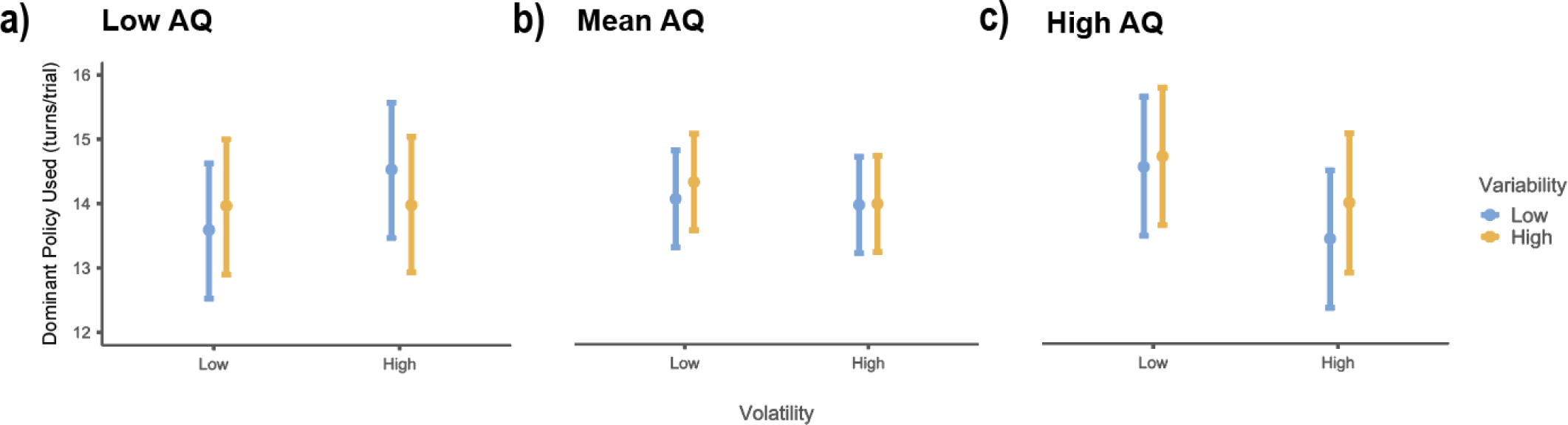
Dominant Policy Use. The turns participants made were categorised into types. This figure shows the number of turns in the participants’ own dominant strategy, controlling for total number of turns. For participants with low AQ (<16, panel a), only for low variability trials (blue), participants used their dominant policy more in high volatility (right) than low (left). For participants with AQ scores within one standard deviation of the mean (panel b), there was no difference between volatility conditions (left/right). In both variability conditions (blue and orange), participants with high AQ (>27, panel c) used their dominant policy more in low volatility (left) than high (right). Error bars are 95% CI.

These findings suggest that different levels of autism traits were associated with differences in the quantity of sampling behaviour, differences in fine-grained movement qualities and differences in the flexibility of policy-selection itself.

### Overall Task Performance: Judgement of Agency

Average accuracy in the judgement of agency task (**Figure 6**) was moderately high across conditions (μ=81.0%, σ=9.12%). MLM results show a significant main effect of variability (F(1,4664)=85.07, p<0.001) such that accuracy was approximately 10% higher in the low variability condition than in the high variability condition. Additionally, there was a significant interaction between variability and volatility (F(1,4665)=8.62, p=0.003). Post-hoc analysis revealed significant differences in all comparisons between the four conditions (z=4.41-8.66, p<0.001) except between low and high volatility when variability remained constant (low/low vs low/high p=0.421, high/low vs high/high p=0.115). This result indicates that while volatility does not make a significant difference to accuracy on its own, the effect of variability on accuracy was stronger under high volatility. There was no significant effect of AQ on accuracy.

**Figure 6.**
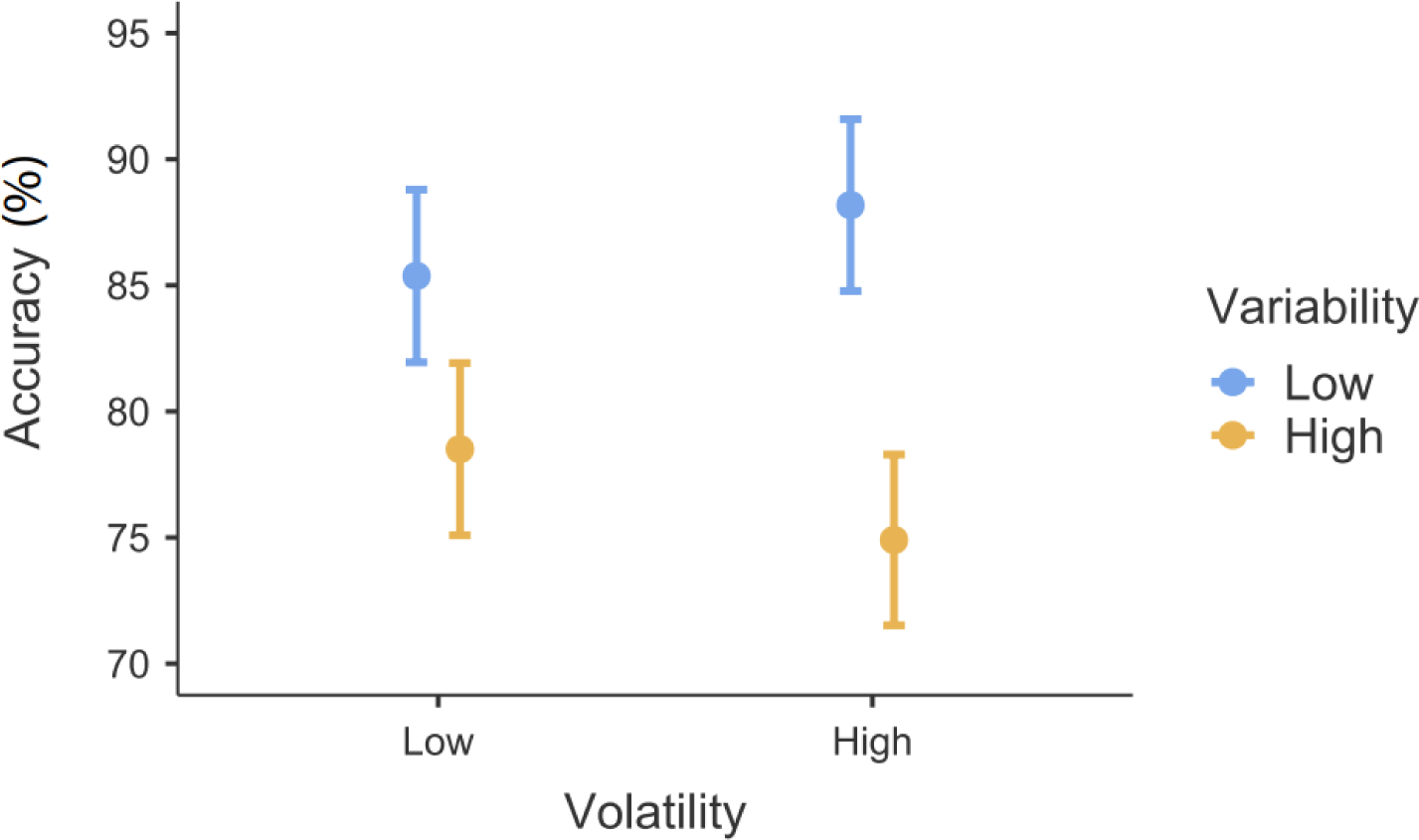
Accuracy. Proportion of trials where participants chose the correct square. Participants were more accurate in low variability (blue) than high variability (orange), and this difference was more pronounced under high volatility (right) than low (left). Error bars are 95% CI.

## Discussion

In this experiment, we closed the action-perception loop to investigate how uncertainty in self-caused sensations influences successive choices about which actions to perform to infer agency. Unlike many previous studies, these actions were freeform and temporally contiguous with ongoing sensory consequences. We showed that action selection changes depending on uncertainty in the mapping between actions and sensory outcomes. We also demonstrate that agency inferences reflect the temporal dynamics of prediction error.

One of the most significant advances of this study on previous designs is the ability to measure and interrogate the temporal dynamics of prediction error, and how this relates to participant behaviour. Using this proxy for prediction error there were particularly interesting findings in the behavioural pattern around hypothesis switches and prediction error minimisation for trials with different judgements of agency. We will now discuss each of these in turn.

Our eye-tracking analysis indicates a hypothesis switch when the participant moves from looking at one square to another and is indicative of a change in the moment to moment beliefs about agency with respect to the candidate square. For action to occur under the active inference account, prediction error comes first, and the action is performed to resolve it. This is consistent with the increasing prediction error leading to a hypothesis switch in our task, indicated by the significant peak in prediction error at the time of the hypothesis switch. The current agential hypothesis is abandoned when the prediction error is too high – there is decreasing evidence that one can achieve one’s expected state with the available actions under the current hypothesis, which leads to a switch that alleviates prediction error. This finding is uniquely consistent with predictive processing (Friston, 2017).

Environmental uncertainty influences this pattern too; after removing trial-wise average prediction error, low variability conditions have a higher prediction error in the hypothesis switch epoch. Also, only in these low variability conditions is there a significant increase from time bin one to time bin two, preceding the switch. Both of these findings suggest that when variability is low, prediction error is allowed to increase for a comparatively longer period of time before the participant decides to switch. This may reflect more reliance on priors in such environments, which allow stable accumulation of evidence for a given hypothesis, and a reluctance to abandon hypotheses in the face of sensory evidence to the contrary.

By looking at the relationship between participants’ agency reports and the trend in prediction error over time, our results suggest that participants could be using these trends to inform their judgment of agency. Agency judgements, whether correct or not, were associated with a more negative prediction error slope. Under the predictive processing account, a correct judgement of agency should be associated with a negative trend in prediction error, and a correct judgement of no-agency should not be associated with prediction error minimisation, as the participant cannot effectively control the stimuli to reduce prediction error. These hypotheses were fully borne out for participants with low AQ scores – when participants correctly judged that they had no agency, the slope of the prediction error was more positive (i.e. failed prediction error minimisation) than when they incorrectly said that they had no agency.

Traditionally, internal representations of agency have been explained using a comparator model. In this model, upon movement, the neural system creates an efference copy of motor commands, which predicts “future states of the motor system and the sensory consequences of movement” (Moore & Obhi, 2012, p. 549). This is then compared with incoming sensory information. In both the comparator and predictive processing accounts, agency is associated with small prediction error, or a match between expected and actual outcomes of actions. The comparator however focuses on net retrospective prediction error and cannot account for hypothesis switches in the face of accumulating prediction error or other changes in future action based on inferences of agency (see also Zaadnoordijk, Besold, and Hunnius (2019)). The predictive processing account positions agency in a broader theory of action and policy selection. So, if the projected reliability of policy-outcome mappings over time under a particular hypothesis (occurrent agency) changes, this account is consistent with a threshold in accumulating prediction error after which the agent switches hypotheses and is especially well equipped if this threshold is sensitive to environmental volatility. Our hypothesis switch ERPE suggests that hypothesis switching is sensitive to volatility when variability is low, with more prediction error around a hypothesis switch when volatility is high.

These results provide a reminder that agents’ ability to discern, and make judgements about, agency arises as they *actively* close the action-perception loop, not just in passive perceptual processes. The results also offer an indication of how agents do this, namely through exploratory titration of prediction error, in a pattern that is sensitive to variability and volatility. It may be that affording agents the opportunity for exploration of the action-perception loop is critical for agency inference and judgement.

Variability also affected the temporal dynamics of prediction error. For instance, under low variability, prediction error was minimised more quickly than in high variability conditions. Comparing the two levels of uncertainty manipulated here, changes to variability caused the most broad-reaching effects. Under high variability, participants were less accurate but spent longer sampling the environment, moved slower but accelerated more quickly, switched hypotheses less frequently and turned less, compared to the low variability conditions. The finding that participants move more under increased variability is consistent with the findings by Wen and Haggard (2020) in a similar judgement of agency paradigm.

While volatility was expected to have effects independent from variability, most of the significant effects for volatility were interactions with variability; volatility only showed two main effects. The first main effect indicated that prediction error was reduced more quickly under high volatility. In our manipulation, the timing of volatile switches was unpredictable, so this effect is likely due to an increased vigilance or sensitivity to incoming information manifesting as an increased learning rate under high volatility (Mathys, Daunizeau, Friston, & Stephan, 2011). The second main effect of volatility indicated that higher volatility was related to higher prediction error in the epoch surrounding hypothesis switches, however this was only true when variability was low. In two further cases, the effect of volatility was only seen in low variability; specifically that participants move faster and switch hypotheses more in high volatility than low. This could reflect an attempt to garner more evidence about the current state of the world before it changes. Lower volatility also increased the effect of variability on frequency of turns and average prediction error across the trial. Conversely, higher volatility increased the effect of variability on task accuracy. Future studies should consider ways of highlighting changes in volatility to enhance the potential effect of higher order uncertainty, such as making them large enough to stand out more saliently to the participant.

It is important to keep in mind too that our analyses of prediction error were limited to a behavioural proxy (combining eye-tracking and mouse movement) for prediction error that does not directly reflect changing internal representations of environmental uncertainty. Future research should consider using neural estimates of prediction error or computational modelling that appropriately changes priors with uncertainty.

Here, we found no difference in accuracy of judgement of agency between healthy participants across a range of autism traits, consistent with previous research comparing autistic and healthy participants on similar measures (David et al., 2008; Grainger et al., 2014; Russell & Hill, 2001; Williams & Happé, 2009; Zalla et al., 2015). As previously noted by Perrykkad and Hohwy (2019) and Zalla and Sperduti (2015), this is in contrast to sense of agency in autism being shown to be reduced under typical experimental paradigms (Sperduti, Pieron, Leboyer, & Zalla, 2014; van Laarhoven, Stekelenburg, Eussen, & Vroomen, 2019). Our study also shows no main effects of AQ on other outcomes, except for a negative association with acceleration.

To our knowledge, Zalla et al. (2015) is the only other case where variability of a similar kind (which they labelled ‘*turbulence’*) was added in a judgement of agency task pertaining to autism, in their case contrasting participants with and without an autism spectrum diagnosis. Their results demonstrated that the accuracy of autistic participants’ agency judgements was less sensitive to differences in variability than the neurotypical group. This study supports our hypothesis that the addition of uncertainty has a distinctive effect on judgement of agency related to autistic traits. While we do not show any significant interactions with AQ in accuracy, our results showed participants with high autism traits were less sensitive to differences in variability in their average prediction error. Since this measure is behavioural, this suggests that participants with high AQ were moving (that is, exploring the environment) in a way that did not reflect underlying differences in variability. Further, AQ was negatively associated with prediction error in the 200ms window surrounding hypothesis switches. This suggests participants with high AQ are switching hypotheses earlier than participants with low AQ, or tolerating less uncertainty before abandoning their current hypothesis (see also Lawson et al. (2017)).

By additionally manipulating volatility, we could demonstrate further effects of uncertainty dependent on AQ. Participants with high AQ were more sensitive to differences in volatility such that only for this group was increased volatility associated with more time spent moving (if only in high variability) and more flexibility in policy selection. This might reflect less consistent or shallower internal models (Perrykkad & Hohwy, 2019), which leads to less precision over all policies in high volatility, and so the selection of one over another fluctuates more frequently. This pattern is the opposite of the low AQ group, where high volatility was associated with more dominant policy use (but only in low variability). This is also consistent with Lawson et al. (2017), who showed that autistic participants update their learning in response to volatility more readily than neurotypical participants.

Of note, our findings with respect to autism are limited to scores on a trait-based measure, which may not generalise to diagnosed autistic populations. Our sample had a high average AQ score compared to what is expected in the general population (Baron-Cohen, Wheelwright, Skinner, Martin, & Clubley, 2001), so our results for “low” AQ may actually be more representative of “average” AQ individuals. While overall the sample size in post-hoc analyses is low, the omnibus interactions were based on modelled trends in the full dataset of continuous AQ scores. Nevertheless, environmental uncertainty might be particularly relevant to action selection for different levels of autistic traits and we do show interactions between uncertainty and AQ. These are worth following up in future studies in diagnosed samples.

In summary, this suggests autistic traits are related to 1) subtle differences in more abstract action policies, which are more sensitive to volatility, 2) smaller differences in prediction error between variability conditions, and 3) a greater propensity to switch hypotheses at a lower prediction error threshold when inferring agency. Notably, despite these differences, there was no significant effect of AQ on overall number of hypothesis switches or on accuracy.

## Conclusion

This experiment shows that uncertainty in the mapping between actions and their outcomes changes not only how effectively participants can identify which stimuli they have control over, but also changes the actions they make and the overall strategies they employ. These changes have downstream impacts on the prediction error which can be used to inform their next action, and their overall response in each trial. In addition, our data illuminates subtle differences in this perception-action loop dependent on autism traits.

## Methods

### Participants

Fifty neurotypical adult participants were recruited. Ten participants were excluded: five participants were removed for technical errors in recording, two for poor quality eye-tracking data (>35% lost trials) and three for poor accuracy (<45%). The final sample of 40 participants were primarily undergraduate students (55%, the remainder had completed tertiary degrees) with an overall mean age of 22.8 years (SD: 3.65, range: 18-34) and included 24 female participants. None of the participants reported neurological conditions, taking medications which affect cognition, nor a history of drug abuse. One participant reported a diagnosis of depression, and one of ADHD, removing these participants did not affect the primary results of interest (see supplementary materials). Two participants reported previously suffering a blow to the head that rendered them unconscious. All participants were fluent in English, had normal or corrected-to-normal vision and 95% were right handed. This study was approved by Monash University Human Research Ethics Committee (Project Number 11396). The experiment was conducted in accordance with the relevant guidelines and regulations, and all participants signed informed consent documents upon commencing the protocol.

### Autism Quotient

None of the participants were previously diagnosed with Autism Spectrum Disorder or its nominal variants. All participants completed the Autism Quotient questionnaire (Baron-Cohen et al., 2001) to quantify autistic traits. The mean AQ score was 21.43 (SD: 5.89, range: 12-38).

### Experimental Task Design and Procedure

For a schematic diagram of the experimental set up, task and experimental manipulation, see **Figure 1** and a video of the task is available at https://figshare.com/s/fd2742b897e21d901dd0 (DOI: 10.26180/5eabbfb9a8aa4).

Testing was conducted in a quiet, darkened room. Participants were seated at a table with a chin rest set to a comfortable height, 84cm from the screen, and approximately 55cm from eye to eye-tracking camera. The task was completed using a computer mouse in the participant’s dominant hand which was hidden in a curtained box (base dimensions: 32cm wide x 30cm deep). Their opposite hand gave judgment of agency responses using the numbers on a keyboard. Participants had self-timed breaks between blocks.

We implemented a variant of *the Squares Task* (Grainger et al., 2014; Russell & Hill, 2001; Williams & Happé, 2009), presented using Psychtoolbox-3.0.14 version beta in Matlab 2017b (Mathworks, Natick, Massachusetts) on a 1920×1080 screen (60Hz refresh rate). Eight randomly coloured squares (100px^2^) appeared in an array at the beginning of each trial. All the squares moved when the mouse was moved and all the squares stopped when the mouse stopped, so participants had to move in order to accurately complete the task. Participants were given 15sec to identify the target square which they controlled. Distracter squares moved at a random angle offset from the vector of mouse movement, and this angle was also independently and randomly changed (and smoothly transitioned) five times in each trial. This means that each distracter square appeared to turn five times when the participant did not initiate a turn, breaking any illusion of control resulting from motor adaptation. There were also less frequent *no-control* trials in which all the eight squares were distracter squares. After the 15sec, all squares froze and were numbered, and prompted an unspeeded numerical response from participants indicating which square they controlled or ‘0’ if they thought they controlled none of the squares.

There were four uncertainty conditions in a 2×2 design (variability x volatility). Some jitter was added to all squares (*variability*), such that depending on the condition, there was a range (95% CI) of random noise around the mean angle input by the mouse (or the distractor offset). This specified range also changed throughout the trial; the number of these changes was specified by the *volatility*. In the *low variability* condition, the distribution switched between a 10° and 30° 95% confidence interval around the mean, and for *high variability*, it switched between 90° and 110°. In the *low volatility* condition, the variability changed three times, while in *high volatility*, there were 10 switches (pseudo-randomly timed with at least 50 frames between). Each trial’s starting distribution was randomly selected. There were two blocks of each condition (variability-volatility pair) with 18 trials per block (16 agentive trials, 2 no-control trials) and block order was randomized for each participant.

Prior to completing the task blocks, participants engaged in an interactive instruction demonstration. Participants then completed a practice block containing sixteen total trials consisting of all trial types, which was excluded from all analyses.

At the end of the experiment there was a short motor control task. In this task, participants were asked to move a perfectly controllable square along a white path as fast and as accurately as possible. There were 10 predesignated paths ranging in length and complexity. This task allowed us to quantify participants’ ability to execute motor intentions.

### Analysis

#### Behaviour

Performance on the motor control task was summarised by multiplying average area traversed outside the white path by average reaction time. This was taken as an index of speed-accuracy trade-off, where low scores indicate better motor performance. Performance on the motor control task did not significantly correlate with AQ (r=0.07, p=0.65) or overall accuracy (r= -0.21, p=0.20), and so was not included as a random effect in any mixed model.

In the ‘squares’ task, *Accurate* trials were those in which participants either correctly identified the target square, or correctly identified that there was no such square (*no-control* trials). Accuracy was the primary measure of overall task performance.

The *time spent moving* on each trial was calculated in seconds. This served as a proxy for environmental sampling, as participants were given freedom to start and stop moving as they pleased though only got task-relevant information by moving.

The *speed* of movement was calculated as the average pixels moved per frame, *acceleration* as change in speed per frame, and *jerk* as the change in acceleration per frame. Derivatives to the level of jerk were analysed to investigate the minimum jerk hypothesis of motor control (Wolpert, 1997) and for its possible relationship to movement trajectories in autism and its traits (Palmer, Paton, Hohwy, & Enticott, 2013; Palmer, Paton, Kirkovski, Enticott, & Hohwy, 2015).

On each frame, the participant’s angle of motion was discretised into one of eight cardinal directions. These were plotted for visual inspection. Participants were found to primarily move in the cardinal directions (up, down, left, right), with smaller peaks at the diagonal midpoints. These plots, in combination with observation of trial replays, informed subsequent policy definition. A *turn* was defined as any change in direction which was preceded by at least three frames of one direction and sustained for at least three frames. More than simply sampling, which also occurred in straight movements, turning involves participant induced intervention on expected stimuli direction. These turns were further grouped into types, which were taken to indicate the participant’s policy. These are pictorially and algorithmically defined in the Supplementary Materials. In brief, six policy types were identified: 1) Horizontal, 2) Vertical, 3) Perpendicular-Cardinal, 4) Non-Cardinal, 5) Hesitant-straight and 6) Circle. Note that rounded corners and circles were redefined for analysis as one turn each as they are taken as a unified intent of intervention by the participant.

While none of these policies has an *a priori* advantage over any other for task performance, we were interested in how flexible each participant was in switching between policies. For each policy, we created a mean percentage of turns that were of that type, across all conditions. We defined a participants’ *dominant policy* as the policy which had the highest percentage of turns across the entire experiment. This allowed us to look at the number of turns in each trial which were of the participants’ dominant policy as compared to alternative policies, as a proxy for exploratory behaviour (i.e., more dominant policy use as exploitative policy selection, less dominant policy use as exploratory). The number of turns on each trial which fell into a participants’ dominant policy were used for this analysis.

#### Eyetracking

Binocular eyetracking data was collected using the SR Research Eyelink 1000 system. For each participant, binocular thirteen-point calibration was conducted; where calibration was unsuccessful for both eyes, one eye was used. The screen x and y coordinates were preprocessed for analysis. Preprocessing involved removing any values outside of the screen bounds, interpolating eyeblinks (as defined by pupil size outside of 1.5 standard deviations below to 2 standard deviations above participant average pupil size), applying a Hanning window of 15 samples (93% overlap) to smooth the eyetrace, and replacing temporarily lost values in one eye with valid data from the other eye (including for whole trials if one eye was excessively noisy). Data was then epoched into trials, and downsampled to match the stimuli framerate for alignment with behavioural data. Trials with poor signal were defined as those with more than 30% of the samples interpolated in both eyes, or whose recorded behavioural data was outside of two standard deviations above or below the participants mean recorded trial length (as the source of these outliers could not be identified). For the final sample of participants, there were a maximum of 65 poor-signal trials (mean = 33.4). Poor-signal trials were removed from all analyses (including behavioural only dependent variables above).

The square the participant was looking at was determined by a novel biased-nearest-object method (see Supplementary Materials). Times of *hypothesis switch* from one square to another were defined as any change in the looked-at square that lasts longer than one frame.

The Euclidean distance between the expected location (had the stimuli followed the mouse) and the actual location of the hypothesised square was calculated as a proxy for *prediction error*. This means that the prediction error is contingent on how quickly the participants move (the error is higher if they move faster) and the quality of their hypothesis. Due to the manipulation, this means that low variability trials accrue less prediction error on average than high variability trials. The *average prediction error* for each participant was calculated across each trial. The *slope of prediction error*, representing the rate of prediction error minimisation, was the slope of the line of best fit of the average prediction error at each time point in each condition for each participant (see **Figures 5a and 5c**). As such, negative values here represent prediction error minimisation, while positive values represent accumulating prediction error.

Finally, given the temporal resolution of our prediction error measure, we were interested in the pattern of prediction error around key temporal events – namely hypothesis switches and changes to the variability distribution (due to volatility). We call these analyses *event-related prediction errors* (ERPE). A one second epoch was centred on the event of interest (time zero) and prediction errors were averaged for each participant in each condition to create an average pattern of activity around the event. Means over five 200ms time bins for each participant were taken for statistical analysis (bin number three is centered on the event onset, see **Figure 6a**). There was no effect of time-bin in the volatility ERPE analysis, hence these are reported in the Supplementary Materials.

### Statistics

All statistical analyses were conducted as Mixed Linear Models (MLM) using Jamovi version 1.1.4 and the GAMLj module (Gallucci, 2019; R Core Team, 2018; The Jamovi Project, 2019). Trial-wise data was used for all dependent variables except prediction error slopes and ERPEs for which condition-wise data was used. Variability and volatility were modelled as simple fixed effect factors, and AQ score was modelled as a continuous fixed effect. All interactions between fixed effects were included. By-participant random intercepts were included to address the non-independence of subject-level observations across trials and capture individual variability in task performance. Compared to traditional methods, this approach affords more sophisticated handling of missing and outlying data, thus improving the accuracy, precision, and generalisability of fixed effect estimates (Singmann & Kellen, 2020). See Table 1 for additional covariates for each model. Degrees of freedom are reported as estimated by the Satterthwaite method. *Post-hoc* analyses were conducted with Bonferroni correction for multiple comparisons and post-hoc p-values are reported with this correction. Post-hoc tests for interactions with AQ were simple effects contrasting participants with low (<Mean-1SD=16, n=6), within one standard deviation from the mean and high (>Mean+1SD=27, n=6) scores for ease of interpretation.

### Data Availability

Dataset used for Results, Table 1, and Figures 2-6 freely available at https://figshare.com/s/77dececaa2b966db4cf7 (DOI:10.26180/5ed0708f103a2).

## Acknowledgements

The authors appreciate the time and effort of their participants without whom this research would not be possible. The authors would also like to thank members of the Cognition and Philosophy Lab, particularly Andrew Corcoran, Jonathan Robinson, Stephen Gadsby, and Andrew McKilliam for their helpful comments on an earlier version of this paper. Thanks also to Chase Sherwell and Bryan Paton for analysis advice.

Perrykkad is supported by an Australian Government Research Training Program (RTP) Scholarship. Hohwy supported this project using Australian Research Council Discovery Grants (DP160102770, DP190101805). Lawson is supported by a Wellcome Trust Henry Dale Fellowship (206691/Z/17/Z) and an Autistica Future Leaders Award (7265). Jamadar is supported by an Australian National Health and Medical Research Council Fellowship (APP1174164) and the Australian Research Council Centre of Excellence for Integrative Brain Function (CE1400100007).

