## Supplementary Material for "The Effect of Uncertainty on Prediction Error in the Action-Perception Loop"

#### Policy Definitions:

##### Horizontal

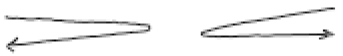

Any turn (surrounded on both sides by at least three frames of direction maintenance) in which the start and end directions are either east or west (right or left).

##### Vertical

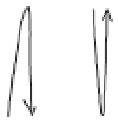

Any turn (surrounded on both sides by at least three frames of direction maintenance) in which the start and end directions are either east or west (right or left).

##### Perpendicular-Cardinal

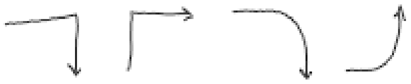

Any turn (surrounded on both sides by at least three frames of direction maintenance) in which the start and end directions are any of the four cardinal directions (right/ left/up/down).

*OR*

Three successive* turns, whose directions move clockwise or anticlockwise but is too short to count as a circle, where the maintenance around the first and last turns are in cardinal directions. This is a *rounded corner* starting and ending in cardinal directions. These get counted as one turn.

##### Non-Cardinal

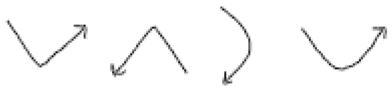

Any turn (surrounded on both sides by at least three frames of direction maintenance) in which the start and end directions are NOT any of the four cardinal directions (diagonals).

*OR*

Three successive* turns, whose directions move clockwise or anticlockwise but is too short to count as a circle, where the maintenance around the first and last turns are NOT in cardinal directions. This is a *rounded corner* starting and ending in diagonals. These get counted as one turn.

##### Hesitant-Straight

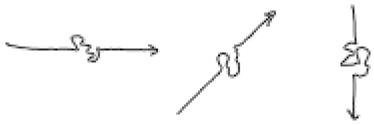

Any time when the direction of travel changes for a little bit, but the directions of the maintenance around the 'hesitation' are in the same direction. Maintenance in any direction in the middle of these cannot be longer than 3 frames (otherwise the turns get counted separately). Can be found in the middle of what is otherwise defined as a rounded turn or a circle also. This counts as a 'turn', because it indicates a decision to maintain direction. Likely occurs frequently as a result of picking up and moving the laser mouse while not intending to change direction at all.

##### Circle

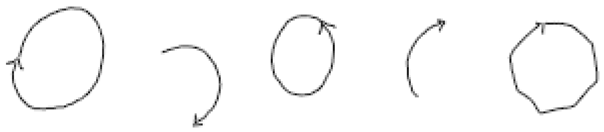

Four successive (not counting adjust-maintains in between) turns whose directions move clockwise or anticlockwise. This counts anything that is at least ¾ of a semi-circle in shape. Each full circle (or any qualifying part of a first circle) gets counted as one turn for this category. Circles are surrounded by at least 3 frames of maintenance in directions non-consistent with a circle, or in the opposite direction of rotation.

*where hesitant-straights in the middle get skipped for counting and labelling this

#### Biased-nearest-object Method for Hypothesis Definition:

For each frame, the shortest Euclidean distance between the eyes and any square was taken to be the square at which the participant was looking, which was taken as a proxy for their *hypothesis* about which square they controlled at that point in time. When two squares were close together (distance to eye positions is less than one squares' length different) and when the nearest square to each eye differed, the hypothesis was biased towards the square which was hypothesised immediately prior. If this was not within the scope of the nearest squares, the nearest square to the average location of the two eyes was chosen.

#### Interaction Results for Movement and Strategy Variables:

Where interactions are not reported here, there were no significant interactions.

##### Speed

There was a significant interaction between variability and volatility (F(1,4661)=16.36, p<0.001). Post-hoc analyses showed that in the low variability/high volatility condition, participants moved faster than the low variability/low volatility (z=3.89, p<0.001), the high variability/high volatility (z=7.18, p<0.001), and the high variability/low volatility conditions (z=5.31, p<0.001). Further, participants moved faster when both variability and volatility were low than when they were both high (z=3.22, p=0.008).

##### Turn Count

There was a significant interaction between variability and volatility (F(1,4661)=4.47, p=0.034). Post-hoc analysis showed the effect of variability was stronger in low volatility conditions (all pairwise contrasts between conditions were significant, z>11.74, p<0.001, except low/low vs low/high and high/low vs. high/high which were non-significant).

##### Hypothesis Switch Count

There was also a significant interaction between variability and volatility (**Figure 5**; $F(1,4661)=6.45$, $p=0.011$). Post-hoc tests showed that in low variability conditions only, participants switched hypotheses more often under high volatility than low volatility ($z=2.79$, $p=0.031$).

#### Volatility Switch ERPE Results

To look at whether changes to the variability distribution due to volatility lead to any differences in prediction error, we performed an MLM on the ERPE centered on volatility switches. In addition to the standard MLM, we included the fixed effect of time-bin, and an additional fixed effect of average prediction error as a covariate. Of the fixed effects of interest, only variability was marginally significant ($F(1,662) = 3.91$, $p=0.05$), such that around the time of volatility switches, high variability trials had greater prediction error than low variability trials ($t(662)=1.98$, $p=0.049$). The lack of interaction with time-bin suggests this difference was not temporally sensitive to the onset of a new variability distribution and similar patterns across uncertainty conditions and autism traits.

#### Effect of Removing Participants with ADHD and Depression:

All models were re-examined with these two participants removed. The following bullet-points report the fixed effects for which the significant effects were altered by removing these two participants.

- For the mixed model using the number of turns per trial as a dependent variable, removing these participants results in a new significant interaction between variability and AQ ($F(1,4441)=4.58$, $p=0.032$). This is down from insignificant $p = 0.074$ in reported sample. Simple effects showed that the difference in variability conditions decreased with increasing AQ score.
- The significant three-way interaction between variability, volatility and AQ for dominant policy use is lost ($F(1,4441)=3.23$, $p=0.072$). Up from $p = 0.039$ in reported sample.
- The marginally significant main effect of volatility for condition-wise prediction error slope is lost ($F(1,108)=3.79$, $p=0.054$).
- The marginally significant main effect of variability for the volatility centered ERPE is lost ($F(1,630)=3.04$, $p=0.082$).

### Full Statistical Models:

For variability and volatility coding, 0 is low, 1 is high.

#### Accuracy Mixed Model

##### Mixed Model

Model Info

| Info | |
|---|---|
| Estimate | Linear mixed model fit by REML |
| Call | Accuracy ~ 1 + Variability + Volatility + AQ + Variability:Volatility + AQ:Volatility + AQ:Variability + AQ:Volatility:Variability+( 1 \| id ) |
| AIC | 4238.694 |
| R-squared Marginal | 0.019 |
| R-squared Conditional | 0.068 |

##### Model Results

Fixed Effect Omnibus tests

| | F | Num df | Den df | p |
|---|---|---|---|---|
| Variability | 85.068 | 1 | 4664.477 | < .001 |
| Volatility | 0.130 | 1 | 4664.483 | 0.718 |
| AQ | 0.057 | 1 | 37.215 | 0.813 |
| Variability ✻ Volatility | 8.617 | 1 | 4664.578 | 0.003 |
| Volatility ✻ AQ | 0.041 | 1 | 4662.239 | 0.840 |
| Variability ✻ AQ | 0.585 | 1 | 4665.167 | 0.445 |
| Variability ✻ Volatility ✻ AQ | 0.682 | 1 | 4664.549 | 0.409 |

Note. Satterthwaite method for degrees of freedom

Fixed Effect Omnibus tests

| | F | Num df | Den df | p |
|---|---|---|---|---|

Fixed Effects Parameter Estimates

| | | | | 95% Confidence Interval | | | | |
|---|---|---|---|---|---|---|---|---|
| Names | Effect | Estimate | SE | Lower | Upper | df | t | p |
| (Intercept) | (Intercept) | 0.817 | 0.015 | 0.789 | 0.846 | 37.382 | 56.026 | < .001 |
| Variability1 | 1 - 0 | -0.101 | 0.011 | -0.122 | -0.079 | 4664.477 | -9.223 | < .001 |
| Volatility1 | 1 - 0 | -0.004 | 0.011 | -0.025 | 0.017 | 4664.483 | -0.361 | 0.718 |
| AQ | AQ | 5.958e-4 | 0.003 | -0.004 | 0.006 | 37.215 | 0.238 | 0.813 |
| Variability1 ✻ Volatility1 | 1 - 0 ✻ 1 - 0 | -0.064 | 0.022 | -0.107 | -0.021 | 4664.578 | -2.935 | 0.003 |
| Volatility1 ✻ AQ | 1 - 0 ✻ AQ | - 3.758e−4 | 0.002 | -0.004 | 0.003 | 4662.239 | -0.202 | 0.840 |
| Variability1 ✻ AQ | 1 - 0 ✻ AQ | 0.001 | 0.002 | -0.002 | 0.005 | 4665.167 | 0.765 | 0.445 |
| Variability1 ✻ Volatility1 ✻ AQ | 1 - 0 ✻ 1 - 0 ✻ AQ | -0.003 | 0.004 | -0.010 | 0.004 | 4664.549 | -0.826 | 0.409 |

Random Components

| Groups | Name | SD | Variance | ICC |
|---|---|---|---|---|
| id | (Intercept) | 0.085 | 0.007 | 0.050 |
| Residual | | 0.374 | 0.140 | |

Fixed Effect Omnibus tests

| | F | Num df | Den df | p |
|---|---|---|---|---|

Note. Number of Obs: 4707 , groups: id , 40

#### Post Hoc Tests

Post Hoc Comparisons - Variability ✻ Volatility

| Comparison | | | | | | | | |
|---|---|---|---|---|---|---|---|---|
| Variability | Volatility | | Variability | Volatility | Difference | SE | z | $p_{bonferroni}$ |
| 0 | 0 | - | 0 | 1 | -0.028 | 0.016 | -1.811 | 0.421 |
| 0 | 0 | - | 1 | 0 | 0.069 | 0.016 | 4.414 | < .001 |
| 0 | 0 | - | 1 | 1 | 0.105 | 0.015 | 6.785 | < .001 |
| 0 | 1 | - | 1 | 1 | 0.133 | 0.015 | 8.661 | < .001 |
| 1 | 0 | - | 0 | 1 | -0.097 | 0.015 | -6.260 | < .001 |
| 1 | 0 | - | 1 | 1 | 0.036 | 0.015 | 2.343 | 0.115 |

### Time Spent Moving Mixed Model

#### Mixed Model

Model Info

| Info | |
|---|---|
| Estimate | Linear mixed model fit by REML |
| Call | TimeSpentMoving ~ 1 + AQ + Volatility + Variability + TimetoMovement + Volatility:Variability + AQ:Variability + AQ:Volatility + AQ:Variability:Volatility+( 1 \| id ) |
| AIC | 13750.596 |
| R-squared Marginal | 0.127 |
| R-squared Conditional | 0.559 |

#### Model Results

Fixed Effect Omnibus tests

| | F | Num df | Den df | p |
|---|---|---|---|---|
| AQ | 0.374 | 1 | 38.003 | 0.544 |
| Volatility | 0.251 | 1 | 4660.287 | 0.616 |
| Variability | 727.707 | 1 | 4660.277 | < .001 |
| TimetoMovement | 431.902 | 1 | 4674.855 | < .001 |
| Volatility ✻ Variability | 2.694 | 1 | 4660.280 | 0.101 |
| AQ ✻ Variability | 2.489 | 1 | 4660.325 | 0.115 |
| AQ ✻ Volatility | 1.702 | 1 | 4660.132 | 0.192 |
| AQ ✻ Volatility ✻ Variability | 11.367 | 1 | 4660.309 | < .001 |

Note. Satterthwaite method for degrees of freedom

Fixed Effects Parameter Estimates

| | | | | 95% Confidence Interval | | | | |
|---|---|---|---|---|---|---|---|---|
| **Names** | **Effect** | **Estimate** | **SE** | **Lower** | **Upper** | **df** | **t** | **p** |
| (Intercept) | (Intercept) | 13.721 | 0.160 | 13.408 | 14.035 | 38.013 | 85.780 | < .001 |
| AQ | AQ | -0.017 | 0.027 | -0.071 | 0.037 | 38.003 | -0.612 | 0.544 |
| Volatility1 | 1 - 0 | -0.015 | 0.030 | -0.073 | 0.043 | 4660.287 | -0.501 | 0.616 |
| Variability1 | 1 - 0 | 0.801 | 0.030 | 0.743 | 0.860 | 4660.277 | 26.976 | < .001 |
| TimetoMovement | TimetoMovement | -0.940 | 0.045 | -1.029 | -0.852 | 4674.855 | -20.782 | < .001 |
| Volatility1 ✻ Variability1 | 1 - 0 ✻ 1 - 0 | -0.098 | 0.059 | -0.214 | 0.019 | 4660.280 | -1.641 | 0.101 |
| AQ ✻ Variability1 | AQ ✻ 1 - 0 | 0.008 | 0.005 | -0.002 | 0.018 | 4660.325 | 1.578 | 0.115 |
| AQ ✻ Volatility1 | AQ ✻ 1 - 0 | -0.007 | 0.005 | -0.017 | 0.003 | 4660.132 | -1.305 | 0.192 |
| AQ ✻ Volatility1 ✻ Variability1 | AQ ✻ 1 - 0 ✻ 1 - 0 | -0.034 | 0.010 | -0.054 | -0.014 | 4660.309 | -3.371 | < .001 |

Random Components

| **Groups** | **Name** | **SD** | **Variance** | **ICC** |
|---|---|---|---|---|
| id | (Intercept) | 1.007 | 1.014 | 0.495 |
| Residual | | 1.017 | 1.035 | |

Note. Number of Obs: 4707 , groups: id , 40

#### Simple Effects

Simple effects of Volatility : Omnibus Tests

| **Moderator levels** | | | | |
|---|---|---|---|---|
| **Variability** | **AQ** | **$X^2$** | **df** | **p** |
| 0 | Mean-1·SD | 0.215 | 1.000 | 0.643 |
| | Mean | 0.643 | 1.000 | 0.423 |
| | Mean+1·SD | 2.594 | 1.000 | 0.107 |
| 1 | Mean-1·SD | 1.617 | 1.000 | 0.204 |
| | Mean | 2.319 | 1.000 | 0.128 |
| | Mean+1·SD | 11.553 | 1.000 | < .001 |

Simple effects of Volatility : Parameter estimates

| **Moderator levels** | | | | | **95% Confidence Interval** | | | |
|---|---|---|---|---|---|---|---|---|
| **Variability** | **AQ** | **contrast** | **Estimate** | **SE** | **Lower** | **Upper** | **z** | **p** |
| 0 | Mean-1·SD | 1 - 0 | -0.028 | 0.060 | -0.145 | 0.089 | -0.463 | 0.643 |
| | Mean | 1 - 0 | 0.034 | 0.042 | -0.049 | 0.117 | 0.802 | 0.423 |
| | Mean+1·SD | 1 - 0 | 0.095 | 0.059 | -0.021 | 0.211 | 1.611 | 0.107 |
| 1 | Mean-1·SD | 1 - 0 | 0.075 | 0.059 | -0.041 | 0.191 | 1.271 | 0.204 |
| | Mean | 1 - 0 | -0.064 | 0.042 | -0.146 | 0.018 | -1.523 | 0.128 |
| | Mean+1·SD | 1 - 0 | -0.203 | 0.060 | -0.319 | -0.086 | -3.399 | < .001 |

Note. Simple effects are estimated keeping constant other independent variable(s) in the model

### Speed Mixed Model

#### Mixed Model

Model Info

| Info | |
|---|---|
| Estimate | Linear mixed model fit by REML |
| Call | MeanSpeed ~ 1 + Variability + Volatility + AQ + Variability:Volatility + Variability:AQ + Volatility:AQ + Variability:Volatility:AQ+( 1 \| id ) |
| AIC | 26650.527 |
| R-squared Marginal | 0.041 |
| R-squared Conditional | 0.641 |

#### Model Results

Fixed Effect Omnibus tests

| | F | Num df | Den df | p |
|---|---|---|---|---|
| Variability | 36.418 | 1 | 4661.161 | < .001 |
| Volatility | 2.196 | 1 | 4661.164 | 0.138 |
| AQ | 2.443 | 1 | 38.004 | 0.126 |
| Variability ✻ Volatility | 16.359 | 1 | 4661.166 | < .001 |
| Variability ✻ AQ | 0.823 | 1 | 4661.186 | 0.364 |
| Volatility ✻ AQ | 0.082 | 1 | 4661.079 | 0.775 |
| Variability ✻ Volatility ✻ AQ | 0.687 | 1 | 4661.165 | 0.407 |

Note. Satterthwaite method for degrees of freedom

Fixed Effects Parameter Estimates

| | | | | 95% Confidence Interval | | | | |
|---|---|---|---|---|---|---|---|---|
| **Names** | **Effect** | **Estimate** | **SE** | **Lower** | **Upper** | **df** | **t** | **p** |
| (Intercept) | (Intercept) | 13.115 | 0.820 | 11.508 | 14.723 | 38.011 | 15.991 | < .001 |
| Variability1 | 1 - 0 | -0.706 | 0.117 | -0.935 | -0.477 | 4661.161 | -6.035 | < .001 |
| Volatility1 | 1 - 0 | 0.173 | 0.117 | -0.056 | 0.403 | 4661.164 | 1.482 | 0.138 |
| AQ | AQ | -0.220 | 0.141 | -0.497 | 0.056 | 38.004 | -1.563 | 0.126 |
| Variability1 ✻ Volatility1 | 1 - 0 ✻ 1 - 0 | -0.947 | 0.234 | -1.405 | -0.488 | 4661.166 | -4.045 | < .001 |
| Variability1 ✻ AQ | 1 - 0 ✻ AQ | -0.018 | 0.020 | -0.057 | 0.021 | 4661.186 | -0.907 | 0.364 |
| Volatility1 ✻ AQ | 1 - 0 ✻ AQ | -0.006 | 0.020 | -0.045 | 0.033 | 4661.079 | -0.286 | 0.775 |
| Variability1 ✻ Volatility1 ✻ AQ | 1 - 0 ✻ 1 - 0 ✻ AQ | 0.033 | 0.040 | -0.045 | 0.111 | 4661.165 | 0.829 | 0.407 |

Random Components

| **Groups** | **Name** | **SD** | **Variance** | **ICC** |
|---|---|---|---|---|
| id | (Intercept) | 5.173 | 26.758 | 0.625 |
| Residual | | 4.007 | 16.054 | |

Note. Number of Obs: 4707 , groups: id , 40

#### Post Hoc Tests

Post Hoc Comparisons - Variability ✻ Volatility

| Comparison | | | | | | | | |
|---|---|---|---|---|---|---|---|---|
| **Variability** | **Volatility** | | **Variability** | **Volatility** | **Difference** | **SE** | **z** | $p_{bonferroni}$ |
| 0 | 0 | - | 0 | 1 | -0.647 | 0.166 | -3.888 | < .001 |

Post Hoc Comparisons - Variability ✻ Volatility

| Comparison | | | | | | | | |
|---|---|---|---|---|---|---|---|---|
| Variability | Volatility | | Variability | Volatility | Difference | SE | z | $p_{bonferroni}$ |
| 0 | 0 | - | 1 | 0 | 0.233 | 0.167 | 1.397 | 0.975 |
| 0 | 0 | - | 1 | 1 | 0.533 | 0.165 | 3.223 | 0.008 |
| 0 | 1 | - | 1 | 1 | 1.179 | 0.164 | 7.180 | < .001 |
| 1 | 0 | - | 0 | 1 | -0.879 | 0.166 | -5.309 | < .001 |
| 1 | 0 | - | 1 | 1 | 0.300 | 0.165 | 1.822 | 0.411 |

Post Hoc Comparisons - Variability

| Comparison | | | | | | |
|---|---|---|---|---|---|---|
| Variability | | Variability | Difference | SE | z | $p_{bonferroni}$ |
| 0 | - | 1 | 0.706 | 0.117 | 6.035 | < .001 |

### Acceleration Mixed Model

#### Mixed Model

Model Info

| Info | |
|---|---|
| Estimate | Linear mixed model fit by REML |
| Call | Acceleration ~ 1 + Variability + Volatility + AQ + Variability:Volatility + Variability:AQ + Volatility:AQ + Variability:Volatility:AQ+( 1 \| id ) |
| AIC | -23077.864 |
| R-squared Marginal | 0.013 |
| R-squared Conditional | 0.077 |

#### Model Results

Fixed Effect Omnibus tests

| | F | Num df | Den df | p |
|---|---|---|---|---|
| Variability | 12.677 | 1 | 4664.138 | < .001 |
| Volatility | 0.779 | 1 | 4664.152 | 0.377 |
| AQ | 5.729 | 1 | 37.799 | 0.022 |
| Variability ✻ Volatility | 1.056 | 1 | 4664.221 | 0.304 |
| Variability ✻ AQ | 0.007 | 1 | 4664.674 | 0.932 |
| Volatility ✻ AQ | 0.379 | 1 | 4662.388 | 0.538 |
| Variability ✻ Volatility ✻ AQ | 0.247 | 1 | 4664.197 | 0.619 |

Note. Satterthwaite method for degrees of freedom

Fixed Effects Parameter Estimates

| | | | | 95% Confidence Interval | | | | |
|---|---|---|---|---|---|---|---|---|
| Names | Effect | Estimate | SE | Lower | Upper | df | t | p |
| (Intercept) | (Intercept) | 0.013 | 8.981e-4 | 0.011 | 0.014 | 37.935 | 14.088 | < .001 |
| Variability1 | 1 - 0 | 0.002 | 5.964e-4 | 9.545e-4 | 0.003 | 4664.138 | 3.560 | < .001 |
| Volatility1 | 1 - 0 | -5.265e−4 | 5.963e-4 | -0.002 | 6.423e-4 | 4664.152 | -0.883 | 0.377 |
| AQ | AQ | -3.691e−4 | 1.542e-4 | -6.713e−4 | -6.684e−5 | 37.799 | -2.393 | 0.022 |
| Variability1 ✻ Volatility1 | 1 - 0 ✻ 1 - 0 | -0.001 | 0.001 | -0.004 | 0.001 | 4664.221 | -1.028 | 0.304 |
| Variability1 ✻ AQ | 1 - 0 ✻ AQ | 8.737e-6 | 1.018e-4 | -1.908e−4 | 2.083e-4 | 4664.674 | 0.086 | 0.932 |
| Volatility1 ✻ AQ | 1 - 0 ✻ AQ | 6.260e-5 | 1.017e-4 | -1.368e−4 | 2.620e-4 | 4662.388 | 0.615 | 0.538 |

Fixed Effects Parameter Estimates

| | | | | 95% Confidence Interval | | | | |
|---|---|---|---|---|---|---|---|---|
| **Names** | **Effect** | **Estimate** | **SE** | **Lower** | **Upper** | **df** | **t** | **p** |
| Variability1 ✻ Volatility1 ✻ AQ | 1 - 0 ✻ 1 - 0 ✻ AQ | -1.012e−4 | 2.036e-4 | -5.003e−4 | 2.978e-4 | 4664.197 | -0.497 | 0.619 |

Random Components

| Groups | Name | SD | Variance | ICC |
|---|---|---|---|---|
| id | (Intercept) | 0.005 | 2.867e-5 | 0.064 |
| Residual | | 0.020 | 4.172e-4 | |

Note. Number of Obs: 4707 , groups: id , 40

#### Post Hoc Tests

Post Hoc Comparisons - Variability

| Comparison | | | | | | |
|---|---|---|---|---|---|---|
| **Variability** | | **Variability** | **Difference** | **SE** | **z** | $p_{bonferroni}$ |
| 0 | - | 1 | -0.002 | 5.964e-4 | -3.560 | < .001 |

#### Post Hoc Tests

Post Hoc Comparisons - Variability

| Comparison | | | | | | |
|---|---|---|---|---|---|---|
| **Variability** | | **Variability** | **Difference** | **SE** | **z** | $p_{bonferroni}$ |
| 0 | - | 1 | -0.0021233 | 5.9637e-4 | -3.5604 | < .001 |

#### Linear Regression

Model Fit Measures

| Model | R | $R^2$ |
|---|---|---|
| 1 | 0.104 | 0.011 |

Model Coefficients - Acceleration

| Predictor | Estimate | SE | t | p |
|---|---|---|---|---|
| Intercept | 0.021 | 0.001 | 17.713 | < .001 |
| AQ | -3.767e−4 | 5.248e-5 | -7.178 | < .001 |

#### Jerk Mixed Model

Model Info

| Info | |
|---|---|
| Estimate | Linear mixed model fit by REML |
| Call | Jerk ~ 1 + Variability + Volatility + AQ + Variability:Volatility + Variability:AQ + Volatility:AQ + Variability:Volatility:AQ+( 1 \| id ) |
| AIC | -32582 |
| R-squared Marginal | 8.5604e-4 |
| R-squared Conditional | 8.5604e-4 |

Note. Results may be uninterpretable or misleading. Try to refine your model.

Note. singular fit

##### Model Results

Fixed Effect Omnibus tests

| | F | Num df | Den df | p |
|---|---|---|---|---|
| Variability | 0.257546 | 1 | 4699.0 | 0.612 |
| Volatility | 0.122248 | 1 | 4699.0 | 0.727 |
| AQ | 0.282417 | 1 | 4699.0 | 0.595 |
| Variability ✻ Volatility | 0.987322 | 1 | 4699.0 | 0.320 |
| Variability ✻ AQ | 1.483438 | 1 | 4699.0 | 0.223 |
| Volatility ✻ AQ | 0.024998 | 1 | 4699.0 | 0.874 |
| Variability ✻ Volatility ✻ AQ | 0.869415 | 1 | 4699.0 | 0.351 |

Note. singular fit

Note. Satterthwaite method for degrees of freedom

Fixed Effects Parameter Estimates

| | | | | 95% Confidence Interval | | | | |
|---|---|---|---|---|---|---|---|---|
| **Names** | **Effect** | **Estimate** | **SE** | **Lower** | **Upper** | **df** | **t** | **p** |
| (Intercept) | (Intercept) | 1.3243e-4 | 1.0928e-4 | -8.1763e–5 | 3.4662e-4 | 4699.0 | 1.21179 | 0.226 |
| Variability1 | 1 - 0 | 1.1092e-4 | 2.1856e-4 | -3.1746e–4 | 5.3930e-4 | 4699.0 | 0.50749 | 0.612 |
| Volatility1 | 1 - 0 | 7.6419e-5 | 2.1856e-4 | -3.5196e–4 | 5.0480e-4 | 4699.0 | 0.34964 | 0.727 |
| AQ | AQ | 9.9132e-6 | 1.8654e-5 | -2.6648e–5 | 4.6474e-5 | 4699.0 | 0.53143 | 0.595 |
| Variability1 ✻ Volatility1 | 1 - 0 ✻ 1 - 0 | 4.3435e-4 | 4.3713e-4 | -4.2241e–4 | 0.0012911 | 4699.0 | 0.99364 | 0.320 |
| Variability1 ✻ AQ | 1 - 0 ✻ AQ | 4.5439e-5 | 3.7308e-5 | -2.7682e–5 | 1.1856e-4 | 4699.0 | 1.21796 | 0.223 |
| Volatility1 ✻ AQ | 1 - 0 ✻ AQ | 5.8986e-6 | 3.7308e-5 | -6.7223e–5 | 7.9020e-5 | 4699.0 | 0.15811 | 0.874 |
| Variability1 ✻ Volatility1 ✻ AQ | 1 - 0 ✻ 1 - 0 ✻ AQ | -6.9573e–5 | 7.4615e-5 | -2.1582e–4 | 7.6670e-5 | 4699.0 | -0.93242 | 0.351 |

Random Components

| **Groups** | **Name** | **SD** | **Variance** | **ICC** |
|---|---|---|---|---|
| id | (Intercept) | 0.0000000 | 0.0000 | 0.0000 |
| Residual | | 0.0074961 | 5.6191e-5 | |

Note. Number of Obs: 4707 , groups: id , 40

### Number of Turns Mixed Model

Model Info

| Info | |
|---|---|
| Estimate | Linear mixed model fit by REML |
| Call | nTurns ~ 1 + Variability + Volatility + AQ + Variability:Volatility + Variability:AQ + Volatility:AQ + Variability:Volatility:AQ+( 1 \| id ) |
| AIC | 35054.526405 |
| R-squared Marginal | 0.036379 |
| R-squared Conditional | 0.532821 |

#### Model Results

Fixed Effect Omnibus tests

| | F | Num df | Den df | p |
|---|---|---|---|---|
| Variability | 346.2197567 | 1 | 4661.184 | < .001 |
| Volatility | 0.0086040 | 1 | 4661.188 | 0.926 |
| AQ | 0.0675365 | 1 | 37.939 | 0.796 |
| Variability ✻ Volatility | 4.4744804 | 1 | 4661.192 | 0.034 |
| Variability ✻ AQ | 3.1868060 | 1 | 4661.224 | 0.074 |
| Volatility ✻ AQ | 0.5633213 | 1 | 4661.055 | 0.453 |
| Variability ✻ Volatility ✻ AQ | 3.7371077 | 1 | 4661.190 | 0.053 |

Fixed Effects Parameter Estimates

| | | | | 95% Confidence Interval | | | | |
|---|---|---|---|---|---|---|---|---|
| Names | Effect | Estimate | SE | Lower | Upper | df | t | p |
| (Intercept) | (Intercept) | 35.435365 | 1.606677 | 32.2863359 | 38.5843951 | 37.949 | 22.055061 | < .001 |

Fixed Effects Parameter Estimates

| | | | | 95% Confidence Interval | | | | |
|---|---|---|---|---|---|---|---|---|
| **Names** | **Effect** | **Estimate** | **SE** | **Lower** | **Upper** | **df** | **t** | **p** |
| Variability 1 | 1 - 0 | -5.333633 | 0.286647 | -5.8954505 | -4.7718154 | 4661.184 | -18.606981 | < .001 |
| Volatility 1 | 1 - 0 | -0.026587 | 0.286632 | -0.5883752 | 0.5352006 | 4661.188 | -0.092758 | 0.926 |
| AQ | AQ | -0.071753 | 0.276104 | -0.6129075 | 0.4694008 | 37.939 | -0.259878 | 0.796 |
| Variability 1 ✻ Volatility 1 | 1 - 0 ✻ 1 - 0 | 1.212699 | 0.573300 | 0.0890523 | 2.3363465 | 4661.192 | 2.115297 | 0.034 |
| Variability 1 ✻ AQ | 1 - 0 ✻ AQ | 0.087372 | 0.048944 | -0.0085554 | 0.1832997 | 4661.224 | 1.785163 | 0.074 |
| Volatility 1 ✻ AQ | 1 - 0 ✻ AQ | 0.036691 | 0.048886 | -0.0591237 | 0.1325065 | 4661.055 | 0.750547 | 0.453 |
| Variability 1 ✻ Volatility 1 ✻ AQ | 1 - 0 ✻ 1 - 0 ✻ AQ | -0.189177 | 0.097859 | -0.3809762 | 0.0026230 | 4661.190 | -1.933160 | 0.053 |

Random Components

| **Groups** | **Name** | **SD** | **Variance** | **ICC** |
|---|---|---|---|---|
| id | (Intercept) | 10.1187 | 102.388 | 0.51518 |
| Residual | | 9.8160 | 96.353 | |

#### Post Hoc Tests

Post Hoc Comparisons - Variability ✻ Volatility

| Comparison | | | | | | | | |
|---|---|---|---|---|---|---|---|---|
| Variability | Volatility | | Variability | Volatility | Difference | SE | z | $p_{bonferroni}$ |
| 0 | 0 | - | 0 | 1 | 0.63294 | 0.40745 | 1.5534 | 0.722 |
| 0 | 0 | - | 1 | 0 | 5.93998 | 0.40833 | 14.5471 | < .001 |
| 0 | 0 | - | 1 | 1 | 5.36022 | 0.40492 | 13.2377 | < .001 |
| 0 | 1 | - | 1 | 1 | 4.72728 | 0.40241 | 11.7473 | < .001 |
| 1 | 0 | - | 0 | 1 | -5.30705 | 0.40582 | -13.0774 | < .001 |
| 1 | 0 | - | 1 | 1 | -0.57976 | 0.40328 | -1.4376 | 0.903 |

Post Hoc Comparisons - Variability

| Comparison | | | | | | |
|---|---|---|---|---|---|---|
| Variability | | Variability | Difference | SE | z | $p_{bonferroni}$ |
| 0 | - | 1 | 5.3336 | 0.28665 | 18.607 | < .001 |

### Number of Dominant Policy Turns Mixed Model

#### Mixed Model

Model Info

| Info | |
|---|---|
| Estimate | Linear mixed model fit by REML |
| Call | nDomPol ~ 1 + nTurns + AQ + Variability + Volatility + Variability:Volatility + Variability:AQ + Volatility:AQ + Variability:Volatility:AQ+( 1 \| id ) |
| AIC | 29580.463 |
| R-squared Marginal | 0.576 |
| R-squared Conditional | 0.633 |

#### Model Results

Fixed Effect Omnibus tests

| | F | Num df | Den df | p |
|---|---|---|---|---|
| nTurns | 3839.669 | 1 | 3842.454 | < .001 |
| AQ | 0.039 | 1 | 36.935 | 0.845 |
| Variability | 0.704 | 1 | 4684.260 | 0.401 |
| Volatility | 1.807 | 1 | 4660.627 | 0.179 |
| Variability ✻ Volatility | 0.574 | 1 | 4660.975 | 0.449 |
| AQ ✻ Variability | 1.853 | 1 | 4661.195 | 0.174 |
| AQ ✻ Volatility | 19.166 | 1 | 4659.774 | < .001 |
| AQ ✻ Variability ✻ Volatility | 4.273 | 1 | 4661.029 | 0.039 |

Note. Satterthwaite method for degrees of freedom

Fixed Effects Parameter Estimates

| | | | | 95% Confidence Interval | | | | |
|---|---|---|---|---|---|---|---|---|
| **Names** | **Effect** | **Estimate** | **SE** | **Lower** | **Upper** | **df** | **t** | **p** |
| (Intercept) | (Intercept) | 14.082 | 0.355 | 13.386 | 14.778 | 36.998 | 39.655 | < .001 |
| nTurns | nTurns | 0.497 | 0.008 | 0.482 | 0.513 | 3842.454 | 61.965 | < .001 |
| AQ | AQ | 0.012 | 0.061 | -0.108 | 0.132 | 36.935 | 0.196 | 0.845 |
| Variability1 | 1 - 0 | 0.140 | 0.167 | -0.187 | 0.467 | 4684.260 | 0.839 | 0.401 |
| Volatility1 | 1 - 0 | -0.217 | 0.161 | -0.533 | 0.099 | 4660.627 | -1.344 | 0.179 |

Fixed Effects Parameter Estimates

| | | | | 95% Confidence Interval | | | | |
|---|---|---|---|---|---|---|---|---|
| **Names** | **Effect** | **Estimate** | **SE** | **Lower** | **Upper** | **df** | **t** | **p** |
| Variability1 ✻ Volatility1 | 1 - 0 ✻ 1 - 0 | -0.244 | 0.323 | -0.876 | 0.388 | 4660.975 | -0.758 | 0.449 |
| AQ ✻ Variability1 | AQ ✻ 1 - 0 | 0.037 | 0.028 | -0.016 | 0.091 | 4661.195 | 1.361 | 0.174 |
| AQ ✻ Volatility1 | AQ ✻ 1 - 0 | -0.120 | 0.027 | -0.174 | -0.066 | 4659.774 | -4.378 | < .001 |
| AQ ✻ Variability1 ✻ Volatility1 | AQ ✻ 1 - 0 ✻ 1 - 0 | 0.114 | 0.055 | 0.006 | 0.222 | 4661.029 | 2.067 | 0.039 |

Random Components

| Groups | Name | SD | Variance | ICC |
|---|---|---|---|---|
| id | (Intercept) | 2.186 | 4.780 | 0.136 |
| Residual | | 5.520 | 30.470 | |

Note. Number of Obs: 4707 , groups: id , 40

#### Simple Effects

Simple effects of Volatility : Omnibus Tests

| Moderator levels | | | |
|---|---|---|---|
| **AQ** | **X²** | **df** | **p** |
| Mean-1·SD | 4.597 | 1.000 | 0.032 |
| Mean | 1.807 | 1.000 | 0.179 |
| Mean+1·SD | 16.367 | 1.000 | < .001 |

Simple effects of Volatility : Omnibus Tests

| Moderator levels | | | |
|---|---|---|---|
| **AQ** | **$X^2$** | **df** | **p** |

Simple effects of Volatility : Parameter estimates

| Moderator levels | | | | 95% Confidence Interval | | | |
|---|---|---|---|---|---|---|---|
| **AQ** | **contrast** | **Estimate** | **SE** | **Lower** | **Upper** | **z** | **p** |
| Mean-1·SD | 1 - 0 | 0.489 | 0.228 | 0.042 | 0.935 | 2.144 | 0.032 |
| Mean | 1 - 0 | -0.217 | 0.161 | -0.533 | 0.099 | -1.344 | 0.179 |
| Mean+1·SD | 1 - 0 | -0.922 | 0.228 | -1.369 | -0.475 | -4.046 | < .001 |

Note. Simple effects are estimated keeping constant other independent variable(s) in the model

### Number of Hypothesis Switches Mixed Model

#### Mixed Model

Model Info

| **Info** | |
|---|---|
| Estimate | Linear mixed model fit by REML |
| Call | nHypSwitches ~ 1 + Variability + Volatility + AQ + Variability:Volatility + AQ:Volatility + AQ:Variability + AQ:Volatility:Variability+( 1 \| id ) |
| AIC | 36948.652 |
| R-squared Marginal | 0.020 |
| R-squared Conditional | 0.578 |

#### Model Results

Fixed Effect Omnibus tests

| | F | Num df | Den df | p |
|---|---|---|---|---|
| Variability | 195.913 | 1 | 4661.203 | < .001 |
| Volatility | 2.045 | 1 | 4661.206 | 0.153 |
| AQ | 0.116 | 1 | 38.005 | 0.736 |
| Variability ✻ Volatility | 6.446 | 1 | 4661.209 | 0.011 |
| Volatility ✻ AQ | 0.143 | 1 | 4661.099 | 0.705 |
| Variability ✻ AQ | 0.280 | 1 | 4661.234 | 0.597 |
| Variability ✻ Volatility ✻ AQ | 0.508 | 1 | 4661.207 | 0.476 |

Note. Satterthwaite method for degrees of freedom

Fixed Effects Parameter Estimates

| | | | | 95% Confidence Interval | | | | |
|---|---|---|---|---|---|---|---|---|
| Names | Effect | Estimate | SE | Lower | Upper | df | t | p |
| (Intercept) | (Intercept) | 42.218 | 2.187 | 37.932 | 46.505 | 38.013 | 19.303 | < .001 |
| Variability1 | 1 - 0 | -4.904 | 0.350 | -5.590 | -4.217 | 4661.203 | -13.997 | < .001 |
| Volatility1 | 1 - 0 | 0.501 | 0.350 | -0.186 | 1.188 | 4661.206 | 1.430 | 0.153 |
| AQ | AQ | -0.128 | 0.376 | -0.865 | 0.609 | 38.005 | -0.340 | 0.736 |
| Variability1 ✻ Volatility1 | 1 - 0 ✻ 1 - 0 | -1.779 | 0.701 | -3.152 | -0.406 | 4661.209 | -2.539 | 0.011 |
| Volatility1 ✻ AQ | 1 - 0 ✻ AQ | -0.023 | 0.060 | -0.140 | 0.094 | 4661.099 | -0.379 | 0.705 |
| Variability1 ✻ AQ | 1 - 0 ✻ AQ | -0.032 | 0.060 | -0.149 | 0.086 | 4661.234 | -0.529 | 0.597 |
| Variability1 ✻ Volatility1 ✻ AQ | 1 - 0 ✻ 1 - 0 ✻ AQ | 0.085 | 0.120 | -0.149 | 0.320 | 4661.207 | 0.712 | 0.476 |

Random Components

| Groups | Name | SD | Variance | ICC |
|---|---|---|---|---|
| id | (Intercept) | 13.785 | 190.038 | 0.569 |
| Residual | | 11.997 | 143.936 | |

Note. Number of Obs: 4707 , groups: id , 40

#### Post Hoc Tests

Post Hoc Comparisons - Variability

| Comparison | | | | | | |
|---|---|---|---|---|---|---|
| **Variability** | | **Variability** | **Difference** | **SE** | **z** | **$p_{bonferroni}$** |
| 0 | - | 1 | 4.904 | 0.350 | 13.997 | < .001 |

Post Hoc Comparisons - Variability ✻ Volatility

| Comparison | | | | | | | | |
|---|---|---|---|---|---|---|---|---|
| **Variability** | **Volatility** | | **Variability** | **Volatility** | **Difference** | **SE** | **z** | **$p_{bonferroni}$** |
| 0 | 0 | - | 0 | 1 | -1.390 | 0.498 | -2.792 | 0.031 |
| 0 | 0 | - | 1 | 0 | 4.014 | 0.499 | 8.044 | < .001 |
| 0 | 0 | - | 1 | 1 | 4.403 | 0.495 | 8.896 | < .001 |
| 0 | 1 | - | 1 | 1 | 5.793 | 0.492 | 11.779 | < .001 |
| 1 | 0 | - | 0 | 1 | -5.405 | 0.496 | -10.897 | < .001 |
| 1 | 0 | - | 1 | 1 | 0.388 | 0.493 | 0.788 | 1.000 |

### Average Prediction Error Mixed Model

#### Mixed Model

Model Info

| Info | |
|---|---|
| Estimate | Linear mixed model fit by REML |
| Call | AvPE ~ 1 + Variability + Volatility + AQ + Variability:Volatility + AQ:Volatility + AQ:Variability + AQ:Volatility:Variability+( 1 \| id ) |
| AIC | 26914.013 |
| R-squared Marginal | 0.085 |
| R-squared Conditional | 0.624 |

#### Model Results

Fixed Effect Omnibus tests

| | F | Num df | Den df | p |
|---|---|---|---|---|
| Variability | 742.628 | 1 | 4661.183 | < .001 |
| Volatility | 1.088 | 1 | 4661.186 | 0.297 |
| AQ | 1.642 | 1 | 38.001 | 0.208 |
| Variability ✻ Volatility | 5.139 | 1 | 4661.189 | 0.023 |
| Volatility ✻ AQ | 0.292 | 1 | 4661.087 | 0.589 |
| Variability ✻ AQ | 31.742 | 1 | 4661.212 | < .001 |
| Variability ✻ Volatility ✻ AQ | 1.026 | 1 | 4661.187 | 0.311 |

Note. Satterthwaite method for degrees of freedom

Fixed Effects Parameter Estimates

| | | | | 95% Confidence Interval | | | | |
|---|---|---|---|---|---|---|---|---|
| **Names** | **Effect** | **Estimate** | **SE** | **Lower** | **Upper** | **df** | **t** | **p** |
| (Intercept) | (Intercept) | 10.436 | 0.783 | 8.901 | 11.970 | 38.008 | 13.327 | < .001 |
| Variability1 | 1 - 0 | 3.281 | 0.120 | 3.045 | 3.517 | 4661.183 | 27.251 | < .001 |
| Volatility1 | 1 - 0 | 0.126 | 0.120 | -0.110 | 0.362 | 4661.186 | 1.043 | 0.297 |
| AQ | AQ | -0.172 | 0.135 | -0.436 | 0.091 | 38.001 | -1.281 | 0.208 |
| Variability1 ✻ Volatility1 | 1 - 0 ✻ 1 - 0 | -0.546 | 0.241 | -1.018 | -0.074 | 4661.189 | -2.267 | 0.023 |
| Volatility1 ✻ AQ | 1 - 0 ✻ AQ | -0.011 | 0.021 | -0.051 | 0.029 | 4661.087 | -0.541 | 0.589 |
| Variability1 ✻ AQ | 1 - 0 ✻ AQ | -0.116 | 0.021 | -0.156 | -0.076 | 4661.212 | -5.634 | < .001 |
| Variability1 ✻ Volatility1 ✻ AQ | 1 - 0 ✻ 1 - 0 ✻ AQ | 0.042 | 0.041 | -0.039 | 0.122 | 4661.187 | 1.013 | 0.311 |

Random Components

| **Groups** | **Name** | **SD** | **Variance** | **ICC** |
|---|---|---|---|---|
| id | (Intercept) | 4.937 | 24.371 | 0.589 |
| Residual | | 4.123 | 17.000 | |

Note. Number of Obs: 4707 , groups: id , 40

#### Post Hoc Tests

Post Hoc Comparisons - Variability ✻ Volatility

| Comparison | | | | | | | | |
|---|---|---|---|---|---|---|---|---|
| **Variability** | **Volatility** | | **Variability** | **Volatility** | **Difference** | **SE** | **z** | $p_{bonferroni}$ |
| 0 | 0 | - | 0 | 1 | -0.399 | 0.171 | -2.329 | 0.119 |
| 0 | 0 | - | 1 | 0 | -3.554 | 0.172 | -20.722 | < .001 |

Post Hoc Comparisons - Variability ✻ Volatility

| Comparison | | | | | | | | |
|---|---|---|---|---|---|---|---|---|
| Variability | Volatility | | Variability | Volatility | Difference | SE | z | $p_{bonferroni}$ |
| 0 | 0 | - | 1 | 1 | -3.407 | 0.170 | -20.030 | < .001 |
| 0 | 1 | - | 1 | 1 | -3.008 | 0.169 | -17.797 | < .001 |
| 1 | 0 | - | 0 | 1 | 3.156 | 0.170 | 18.512 | < .001 |
| 1 | 0 | - | 1 | 1 | 0.147 | 0.169 | 0.870 | 1.000 |

#### Simple Effects

Simple effects of Variability : Omnibus Tests

| Moderator levels | | | |
|---|---|---|---|
| AQ | $X^2$ | df | p |
| Mean-1·SD | 540.731 | 1.000 | < .001 |
| Mean | 742.628 | 1.000 | < .001 |
| Mean+1·SD | 233.394 | 1.000 | < .001 |

Simple effects of Variability : Parameter estimates

| Moderator levels | | | | 95% Confidence Interval | | | |
|---|---|---|---|---|---|---|---|
| AQ | contrast | Estimate | SE | Lower | Upper | z | p |
| Mean-1·SD | 1 - 0 | 3.960 | 0.170 | 3.626 | 4.294 | 23.254 | < .001 |
| Mean | 1 - 0 | 3.281 | 0.120 | 3.045 | 3.517 | 27.251 | < .001 |
| Mean+1·SD | 1 - 0 | 2.602 | 0.170 | 2.269 | 2.936 | 15.277 | < .001 |

Note. Simple effects are estimated keeping constant other independent variable(s) in the model

#### Condition-wise Prediction Error Slope Mixed Model

#### Mixed Model

Model Info

| Info | |
|---|---|
| Estimate | Linear mixed model fit by REML |
| Call | ConditionAvPEGradient ~ 1 + Variability + Volatility + AQ + Variability:Volatility + Variability:AQ + Volatility:AQ + Variability:Volatility:AQ+( 1 \| id ) |
| AIC | -1131.354 |
| R-squared Marginal | 0.104 |
| R-squared Conditional | 0.749 |

#### Model Results

Fixed Effect Omnibus tests

| | F | Num df | Den df | p |
|---|---|---|---|---|
| Variability | 58.154 | 1 | 114.000 | < .001 |
| Volatility | 3.959 | 1 | 114.000 | 0.049 |
| AQ | 0.260 | 1 | 38.000 | 0.613 |
| Variability ✻ Volatility | 0.217 | 1 | 114.000 | 0.643 |
| Variability ✻ AQ | 0.779 | 1 | 114.000 | 0.379 |
| Volatility ✻ AQ | 0.052 | 1 | 114.000 | 0.819 |
| Variability ✻ Volatility ✻ AQ | 0.020 | 1 | 114.000 | 0.887 |

Note. Satterthwaite method for degrees of freedom

Fixed Effects Parameter Estimates

| | | | | 95% Confidence Interval | | | | |
|---|---|---|---|---|---|---|---|---|
| **Names** | **Effect** | **Estimate** | **SE** | **Lower** | **Upper** | **df** | **t** | **p** |
| (Intercept) | (Intercept) | 4.350e-4 | 9.956e-4 | -0.002 | 0.002 | 38.000 | 0.437 | 0.665 |
| Variability1 | 1 - 0 | 0.005 | 5.923e-4 | 0.003 | 0.006 | 114.000 | 7.626 | < .001 |
| Volatility1 | 1 - 0 | -0.001 | 5.923e-4 | -0.002 | -1.762e−5 | 114.000 | -1.990 | 0.049 |
| AQ | AQ | 8.719e-5 | 1.711e-4 | -2.482e−4 | 4.226e-4 | 38.000 | 0.509 | 0.613 |
| Variability1 ✻ Volatility1 | 1 - 0 ✻ 1 - 0 | 5.514e-4 | 0.001 | -0.002 | 0.003 | 114.000 | 0.465 | 0.643 |
| Variability1 ✻ AQ | 1 - 0 ✻ AQ | -8.987e−5 | 1.018e-4 | -2.894e−4 | 1.097e-4 | 114.000 | -0.883 | 0.379 |
| Volatility1 ✻ AQ | 1 - 0 ✻ AQ | -2.329e−5 | 1.018e-4 | -2.228e−4 | 1.763e-4 | 114.000 | -0.229 | 0.819 |
| Variability1 ✻ Volatility1 ✻ AQ | 1 - 0 ✻ 1 - 0 ✻ AQ | -2.900e−5 | 2.036e-4 | -4.281e−4 | 3.701e-4 | 114.000 | -0.142 | 0.887 |

Random Components

| **Groups** | **Name** | **SD** | **Variance** | **ICC** |
|---|---|---|---|---|
| id | (Intercept) | 0.006 | 3.614e-5 | 0.720 |
| Residual | | 0.004 | 1.403e-5 | |

Note. Number of Obs: 160 , groups: id , 40

#### Post Hoc Tests

Post Hoc Comparisons - Variability

| Comparison | | | | | | | |
|---|---|---|---|---|---|---|---|
| Variability | | Variability | Difference | SE | t | df | $p_{bonferroni}$ |
| 0 | - | 1 | -0.005 | 5.923e-4 | -7.626 | 114.000 | < .001 |

Post Hoc Comparisons - Volatility

| Comparison | | | | | | | |
|---|---|---|---|---|---|---|---|
| Volatility | | Volatility | Difference | SE | t | df | $p_{bonferroni}$ |
| 0 | - | 1 | 0.001 | 5.923e-4 | 1.990 | 114.000 | 0.049 |

### Prediction Error Slope by Accuracy and Agency Mixed Model

#### Mixed Model

Model Info

| Info | |
|---|---|
| Estimate | Linear mixed model fit by REML |
| Call | AgAccAvPEGradient ~ 1 + JudgedAgency + Accuracy + AQ + JudgedAgency:Accuracy + Accuracy:AQ + JudgedAgency:AQ + Accuracy:JudgedAgency:AQ+( 1 \| id ) |
| AIC | -1083.147 |
| R-squared Marginal | 0.168 |
| R-squared Conditional | 0.746 |

#### Model Results

Fixed Effect Omnibus tests

| | F | Num df | Den df | p |
|---|---|---|---|---|
| JudgedAgency | 82.886 | 1 | 113.120 | < .001 |
| Accuracy | 1.279 | 1 | 113.120 | 0.260 |
| AQ | 0.001 | 1 | 38.007 | 0.971 |
| JudgedAgency ✻ Accuracy | 12.785 | 1 | 113.120 | < .001 |
| Accuracy ✻ AQ | 0.512 | 1 | 113.071 | 0.476 |
| JudgedAgency ✻ AQ | 0.398 | 1 | 113.071 | 0.529 |
| JudgedAgency ✻ Accuracy ✻ AQ | 5.684 | 1 | 113.071 | 0.019 |

Note. Satterthwaite method for degrees of freedom

Fixed Effects Parameter Estimates

| | | | | 95% Confidence Interval | | | | |
|---|---|---|---|---|---|---|---|---|
| **Names** | **Effect** | **Estimate** | **SE** | **Lower** | **Upper** | **df** | **t** | **p** |
| (Intercept) | (Intercept) | 0.004 | 0.001 | 0.001 | 0.006 | 38.057 | 3.285 | 0.002 |
| JudgedAgency1 | 1 - 0 | -0.006 | 6.892e-4 | -0.008 | -0.005 | 113.120 | -9.104 | < .001 |
| Accuracy1 | 1 - 0 | -7.794e−4 | 6.892e-4 | -0.002 | 5.714e-4 | 113.120 | -1.131 | 0.260 |
| AQ | AQ | -6.890e−6 | 1.873e-4 | -3.740e−4 | 3.602e-4 | 38.007 | -0.037 | 0.971 |
| JudgedAgency1 ✻ Accuracy1 | 1 - 0 ✻ 1 - 0 | -0.005 | 0.001 | -0.008 | -0.002 | 113.120 | -3.576 | < .001 |
| Accuracy1 ✻ AQ | 1 - 0 ✻ AQ | -8.447e−5 | 1.181e-4 | -3.159e−4 | 1.469e-4 | 113.071 | -0.715 | 0.476 |
| JudgedAgency1 ✻ AQ | 1 - 0 ✻ AQ | 7.449e-5 | 1.181e-4 | -1.569e−4 | 3.059e-4 | 113.071 | 0.631 | 0.529 |

Fixed Effects Parameter Estimates

| | | | | 95% Confidence Interval | | | | |
|---|---|---|---|---|---|---|---|---|
| **Names** | **Effect** | **Estimate** | **SE** | **Lower** | **Upper** | **df** | **t** | **p** |
| JudgedAgency1 ✻ Accuracy1 ✻ AQ | 1 - 0 ✻ 1 - 0 ✻ AQ | 5.630e-4 | 2.361e-4 | 1.002e-4 | 0.001 | 113.071 | 2.384 | 0.019 |

Random Components

| Groups | Name | SD | Variance | ICC |
|---|---|---|---|---|
| id | (Intercept) | 0.007 | 4.277e-5 | 0.694 |
| Residual | | 0.004 | 1.884e-5 | |

Note. Number of Obs: 159 , groups: id , 40

#### Post Hoc Tests

Post Hoc Comparisons - JudgedAgency ✻ Accuracy

| Comparison | | | | | | | | | |
|---|---|---|---|---|---|---|---|---|---|
| **JudgedAgency** | **Accuracy** | | **JudgedAgency** | **Accuracy** | **Difference** | **SE** | **t** | **df** | **$p_{bonferroni}$** |
| 0 | 0 | - | 0 | 1 | -0.002 | 9.707e-4 | -1.736 | 113.001 | 0.512 |
| 0 | 0 | - | 1 | 0 | 0.004 | 9.787e-4 | 3.893 | 113.119 | 0.001 |
| 0 | 0 | - | 1 | 1 | 0.007 | 9.707e-4 | 7.267 | 113.001 | < .001 |
| 0 | 1 | - | 1 | 1 | 0.009 | 9.707e-4 | 9.003 | 113.001 | < .001 |
| 1 | 0 | - | 0 | 1 | -0.005 | 9.787e-4 | -5.615 | 113.119 | < .001 |

Post Hoc Comparisons - JudgedAgency ✻ Accuracy

| Comparison | | | | | | | | | |
|---|---|---|---|---|---|---|---|---|---|
| JudgedAgency | Accuracy | | JudgedAgency | Accuracy | Difference | SE | t | df | $p_{bonferroni}$ |
| 1 | 0 | - | 1 | 1 | 0.003 | 9.787e-4 | 3.314 | 113.119 | 0.007 |

Post Hoc Comparisons - JudgedAgency

| Comparison | | | | | | | |
|---|---|---|---|---|---|---|---|
| JudgedAgency | | JudgedAgency | Difference | SE | t | df | $p_{bonferroni}$ |
| 0 | - | 1 | 0.006 | 6.892e-4 | 9.104 | 113.061 | < .001 |

#### Simple Effects

Simple effects of Accuracy : Omnibus Tests

| Moderator levels | | | | | |
|---|---|---|---|---|---|
| JudgedAgency | AQ | F | Num df | Den df | p |
| 0 | Mean-1·SD | 7.744 | 1.000 | 113.060 | 0.006 |
| | Mean | 3.013 | 1.000 | 113.060 | 0.085 |
| | Mean+1·SD | 0.110 | 1.000 | 113.060 | 0.741 |
| 1 | Mean-1·SD | 10.058 | 1.000 | 113.180 | 0.002 |
| | Mean | 10.985 | 1.000 | 113.180 | 0.001 |
| | Mean+1·SD | 2.294 | 1.000 | 113.080 | 0.133 |

Simple effects of Accuracy : Parameter estimates

| Moderator levels | 95% Confidence Interval |
|---|---|

Simple effects of Accuracy : Omnibus Tests

| Moderator levels | | | | | |
|---|---|---|---|---|---|
| **JudgedAgency** | **AQ** | **F** | **Num df** | **Den df** | **p** |

| JudgedAgency | AQ | contrast | Estimate | SE | Lower | Upper | df | t | p |
|---|---|---|---|---|---|---|---|---|---|
| 0 | Mean-1·SD | 1 - 0 | 0.004 | 0.001 | 0.001 | 0.007 | 113.060 | 2.783 | 0.006 |
| | Mean | 1 - 0 | 0.002 | 9.707e-4 | -2.382e−4 | 0.004 | 113.060 | 1.736 | 0.085 |
| | Mean+1·SD | 1 - 0 | -4.563e−4 | 0.001 | -0.003 | 0.002 | 113.060 | -0.331 | 0.741 |
| 1 | Mean-1·SD | 1 - 0 | -0.004 | 0.001 | -0.007 | -0.002 | 113.179 | -3.171 | 0.002 |
| | Mean | 1 - 0 | -0.003 | 9.787e-4 | -0.005 | -0.001 | 113.178 | -3.314 | 0.001 |
| | Mean+1·SD | 1 - 0 | -0.002 | 0.001 | -0.005 | 6.439e-4 | 113.080 | -1.515 | 0.133 |

#### Estimated Marginal Means

JudgedAgency

| | | | | 95% Confidence Interval | |
|---|---|---|---|---|---|
| **JudgedAgency** | **Mean** | **SE** | **df** | **Lower** | **Upper** |
| 0 | 0.007 | 0.001 | 45.757 | 0.004 | 0.009 |
| 1 | 4.434e-4 | 0.001 | 46.012 | -0.002 | 0.003 |

Note. Estimated means are estimated averaging across interacting variables

JudgedAgency

| | | | | 95% Confidence Interval | |
|---|---|---|---|---|---|
| **JudgedAgency** | **Mean** | **SE** | **df** | **Lower** | **Upper** |

Accuracy

| | | | | 95% Confidence Interval | |
|---|---|---|---|---|---|
| **Accuracy** | **Mean** | **SE** | **df** | **Lower** | **Upper** |
| 0 | 0.004 | 0.001 | 46.012 | 0.002 | 0.006 |
| 1 | 0.003 | 0.001 | 45.757 | 8.915e-4 | 0.005 |

Note. Estimated means are estimated averaging across interacting variables

Accuracy:JudgedAgency

| | | | | | 95% Confidence Interval | |
|---|---|---|---|---|---|---|
| **Accuracy** | **JudgedAgency** | **Mean** | **SE** | **df** | **Lower** | **Upper** |
| 0 | 0 | 0.006 | 0.001 | 62.125 | 0.003 | 0.008 |
| 1 | 0 | 0.008 | 0.001 | 62.125 | 0.005 | 0.010 |
| 0 | 1 | 0.002 | 0.001 | 63.180 | -4.272e−4 | 0.005 |

JudgedAgency

| | | | | 95% Confidence Interval | | |
|---|---|---|---|---|---|---|
| **JudgedAgency** | | **Mean** | **SE** | **df** | **Lower** | **Upper** |
| 1 | 1 | -0.001 | 0.001 | 62.125 | -0.004 | 0.001 |

Note. Estimated means are estimated keeping constant other independent variable(s) in the model to the mean

### Volatility ERPE Mixed Model

#### Mixed Model

Model Info

| **Info** | |
|---|---|
| Estimate | Linear mixed model fit by REML |
| Call | VolSwitchERPE ~ 1 + avPE + Volatility + Variability + AQ + TimeBin + Volatility:Variability + Volatility:TimeBin + Variability:TimeBin + AQ:TimeBin + AQ:Variability + AQ:Volatility + Volatility:Variability:TimeBin + AQ:TimeBin:Variability + AQ:TimeBin:Volatility + AQ:Variability:Volatility + AQ:TimeBin:Variability:Volatility+( 1 \| id ) |
| AIC | 2504.522 |
| R-squared Marginal | 0.962 |
| R-squared Conditional | 0.964 |

#### Model Results

Fixed Effect Omnibus tests

| | **F** | **Num df** | **Den df** | **p** |
|---|---|---|---|---|
| avPE | 10716.298 | 1 | 51.346 | < .001 |
| Volatility | 0.520 | 1 | 722.053 | 0.471 |
| Variability | 3.909 | 1 | 662.430 | 0.048 |

Fixed Effect Omnibus tests

| | F | Num df | Den df | p |
|---|---|---|---|---|
| AQ | 0.286 | 1 | 37.521 | 0.596 |
| TimeBin | 0.155 | 4 | 721.814 | 0.961 |
| Volatility ✻ Variability | 0.406 | 1 | 722.935 | 0.524 |
| Volatility ✻ TimeBin | 1.000 | 4 | 721.814 | 0.407 |
| Variability ✻ TimeBin | 0.293 | 4 | 721.814 | 0.883 |
| AQ ✻ TimeBin | 0.067 | 4 | 721.814 | 0.992 |
| Variability ✻ AQ | 0.516 | 1 | 729.816 | 0.473 |
| Volatility ✻ AQ | 2.289 | 1 | 721.995 | 0.131 |
| Volatility ✻ Variability ✻ TimeBin | 0.572 | 4 | 721.814 | 0.683 |
| Variability ✻ AQ ✻ TimeBin | 0.533 | 4 | 721.814 | 0.712 |
| Volatility ✻ AQ ✻ TimeBin | 0.360 | 4 | 721.814 | 0.837 |
| Volatility ✻ Variability ✻ AQ | 0.157 | 1 | 722.043 | 0.692 |
| Volatility ✻ Variability ✻ AQ ✻ TimeBin | 0.095 | 4 | 721.814 | 0.984 |

Fixed Effects Parameter Estimates

| | | | | 95% Confidence Interval | | | | |
|---|---|---|---|---|---|---|---|---|
| **Names** | **Effect** | **Estimate** | **SE** | **Lower** | **Upper** | **df** | **t** | **p** |
| (Intercept) | (Intercept) | 10.726 | 0.051 | 10.626 | 10.826 | 37.115 | 210.753 | < .001 |
| avPE | avPE | 1.010 | 0.010 | 0.991 | 1.029 | 51.346 | 103.520 | < .001 |
| Volatility1 | 2 - 1 | 0.054 | 0.076 | -0.094 | 0.203 | 722.053 | 0.721 | 0.471 |
| Variability1 | 2 - 1 | 0.162 | 0.082 | 0.001 | 0.324 | 662.430 | 1.977 | 0.048 |
| AQ | AQ | -0.005 | 0.009 | -0.022 | 0.013 | 37.521 | -0.535 | 0.596 |
| TimeBin1 | 2 - 1 | 0.025 | 0.119 | -0.209 | 0.259 | 721.814 | 0.209 | 0.834 |
| TimeBin2 | 3 - 1 | 0.017 | 0.119 | -0.217 | 0.251 | 721.814 | 0.142 | 0.887 |

Fixed Effects Parameter Estimates

| | | | | 95% Confidence Interval | | | | |
|---|---|---|---|---|---|---|---|---|
| Names | Effect | Estimate | SE | Lower | Upper | df | t | p |
| TimeBin3 | 4 - 1 | -0.046 | 0.119 | -0.280 | 0.188 | 721.814 | -0.387 | 0.699 |
| TimeBin4 | 5 - 1 | 0.041 | 0.119 | -0.194 | 0.275 | 721.814 | 0.340 | 0.734 |
| Volatility1 ✻ Variability1 | 2 - 1 ✻ 2 - 1 | -0.096 | 0.151 | -0.393 | 0.200 | 722.935 | -0.637 | 0.524 |
| Volatility1 ✻ TimeBin1 | 2 - 1 ✻ 2 - 1 | 0.028 | 0.239 | -0.440 | 0.496 | 721.814 | 0.116 | 0.907 |
| Volatility1 ✻ TimeBin2 | 2 - 1 ✻ 3 - 1 | -0.265 | 0.239 | -0.734 | 0.203 | 721.814 | -1.110 | 0.267 |
| Volatility1 ✻ TimeBin3 | 2 - 1 ✻ 4 - 1 | -0.348 | 0.239 | -0.816 | 0.120 | 721.814 | -1.456 | 0.146 |
| Volatility1 ✻ TimeBin4 | 2 - 1 ✻ 5 - 1 | -0.048 | 0.239 | -0.517 | 0.420 | 721.814 | -0.202 | 0.840 |
| Variability1 ✻ TimeBin1 | 2 - 1 ✻ 2 - 1 | -0.013 | 0.239 | -0.481 | 0.455 | 721.814 | -0.054 | 0.957 |
| Variability1 ✻ TimeBin2 | 2 - 1 ✻ 3 - 1 | 0.203 | 0.239 | -0.265 | 0.671 | 721.814 | 0.850 | 0.395 |
| Variability1 ✻ TimeBin3 | 2 - 1 ✻ 4 - 1 | 0.103 | 0.239 | -0.366 | 0.571 | 721.814 | 0.430 | 0.667 |
| Variability1 ✻ TimeBin4 | 2 - 1 ✻ 5 - 1 | 0.013 | 0.239 | -0.455 | 0.482 | 721.814 | 0.056 | 0.956 |
| AQ ✻ TimeBin1 | AQ ✻ 2 - 1 | -0.007 | 0.021 | -0.048 | 0.033 | 721.814 | -0.362 | 0.718 |
| AQ ✻ TimeBin2 | AQ ✻ 3 - 1 | -0.008 | 0.021 | -0.049 | 0.032 | 721.814 | -0.408 | 0.683 |
| AQ ✻ TimeBin3 | AQ ✻ 4 - 1 | -0.005 | 0.021 | -0.045 | 0.036 | 721.814 | -0.223 | 0.824 |
| AQ ✻ TimeBin4 | AQ ✻ 5 - 1 | -0.009 | 0.021 | -0.049 | 0.031 | 721.814 | -0.450 | 0.653 |
| Variability1 ✻ AQ | 2 - 1 ✻ AQ | -0.009 | 0.013 | -0.035 | 0.016 | 729.816 | -0.718 | 0.473 |
| Volatility1 ✻ AQ | 2 - 1 ✻ AQ | 0.020 | 0.013 | -0.006 | 0.045 | 721.995 | 1.513 | 0.131 |

Fixed Effects Parameter Estimates

| | | | | 95% Confidence Interval | | | | |
|---|---|---|---|---|---|---|---|---|
| **Names** | **Effect** | **Estimate** | **SE** | **Lower** | **Upper** | **df** | **t** | **p** |
| Volatility1 ✻ Variability1 ✻ TimeBin1 | 2 - 1 ✻ 2 - 1 ✻ 2 - 1 | 0.356 | 0.478 | -0.581 | 1.293 | 721.814 | 0.745 | 0.456 |
| Volatility1 ✻ Variability1 ✻ TimeBin2 | 2 - 1 ✻ 2 - 1 ✻ 3 - 1 | 0.134 | 0.478 | -0.802 | 1.071 | 721.814 | 0.281 | 0.779 |
| Volatility1 ✻ Variability1 ✻ TimeBin3 | 2 - 1 ✻ 2 - 1 ✻ 4 - 1 | 0.075 | 0.478 | -0.861 | 1.012 | 721.814 | 0.158 | 0.875 |
| Volatility1 ✻ Variability1 ✻ TimeBin4 | 2 - 1 ✻ 2 - 1 ✻ 5 - 1 | 0.630 | 0.478 | -0.307 | 1.566 | 721.814 | 1.318 | 0.188 |
| Variability1 ✻ AQ ✻ TimeBin1 | 2 - 1 ✻ AQ ✻ 2 - 1 | -0.044 | 0.041 | -0.124 | 0.036 | 721.814 | -1.071 | 0.284 |
| Variability1 ✻ AQ ✻ TimeBin2 | 2 - 1 ✻ AQ ✻ 3 - 1 | -0.005 | 0.041 | -0.085 | 0.076 | 721.814 | -0.114 | 0.909 |
| Variability1 ✻ AQ ✻ TimeBin3 | 2 - 1 ✻ AQ ✻ 4 - 1 | 0.012 | 0.041 | -0.069 | 0.092 | 721.814 | 0.291 | 0.771 |
| Variability1 ✻ AQ ✻ TimeBin4 | 2 - 1 ✻ AQ ✻ 5 - 1 | -0.016 | 0.041 | -0.096 | 0.065 | 721.814 | -0.381 | 0.703 |
| Volatility1 ✻ AQ ✻ TimeBin1 | 2 - 1 ✻ AQ ✻ 2 - 1 | 0.007 | 0.041 | -0.073 | 0.088 | 721.814 | 0.171 | 0.865 |
| Volatility1 ✻ AQ ✻ TimeBin2 | 2 - 1 ✻ AQ ✻ 3 - 1 | 0.031 | 0.041 | -0.050 | 0.111 | 721.814 | 0.753 | 0.452 |
| Volatility1 ✻ AQ ✻ TimeBin3 | 2 - 1 ✻ AQ ✻ 4 - 1 | 0.040 | 0.041 | -0.040 | 0.121 | 721.814 | 0.979 | 0.328 |
| Volatility1 ✻ AQ ✻ TimeBin4 | 2 - 1 ✻ AQ ✻ 5 - 1 | 0.032 | 0.041 | -0.049 | 0.112 | 721.814 | 0.771 | 0.441 |
| Volatility1 ✻ Variability1 ✻ AQ | 2 - 1 ✻ 2 - 1 ✻ AQ | 0.010 | 0.026 | -0.041 | 0.061 | 722.043 | 0.396 | 0.692 |

Fixed Effects Parameter Estimates

| | | | | 95% Confidence Interval | | | | |
|---|---|---|---|---|---|---|---|---|
| **Names** | **Effect** | **Estimate** | **SE** | **Lower** | **Upper** | **df** | **t** | **p** |
| Volatility1 ✻ Variability1 ✻ AQ ✻ TimeBin1 | 2 - 1 ✻ 2 - 1 ✻ AQ ✻ 2 - 1 | 0.034 | 0.082 | -0.127 | 0.195 | 721.814 | 0.418 | 0.676 |
| Volatility1 ✻ Variability1 ✻ AQ ✻ TimeBin2 | 2 - 1 ✻ 2 - 1 ✻ AQ ✻ 3 - 1 | 0.003 | 0.082 | -0.158 | 0.164 | 721.814 | 0.037 | 0.970 |
| Volatility1 ✻ Variability1 ✻ AQ ✻ TimeBin3 | 2 - 1 ✻ 2 - 1 ✻ AQ ✻ 4 - 1 | -0.012 | 0.082 | -0.173 | 0.149 | 721.814 | -0.149 | 0.882 |
| Volatility1 ✻ Variability1 ✻ AQ ✻ TimeBin4 | 2 - 1 ✻ 2 - 1 ✻ AQ ✻ 5 - 1 | 0.017 | 0.082 | -0.144 | 0.178 | 721.814 | 0.212 | 0.832 |

Random Components

| **Groups** | **Name** | **SD** | **Variance** | **ICC** |
|---|---|---|---|---|
| id | (Intercept) | 0.216 | 0.047 | 0.039 |
| Residual | | 1.068 | 1.142 | |

#### Post Hoc Tests

Post Hoc Comparisons - Variability

| **Comparison** | | | | | | | |
|---|---|---|---|---|---|---|---|
| **Variability** | | **Variability** | **Difference** | **SE** | **t** | **df** | **$p_{bonferroni}$** |
| 1 | - | 2 | -0.162 | 0.082 | -1.975 | 662.208 | 0.049 |

### Hypothesis Switch ERPE Mixed Model

#### Mixed Model

Model Info

| Info | |
|---|---|
| Estimate | Linear mixed model fit by REML |
| Call | HypSwitchERPE ~ 1 + Variability + Volatility + TimeBin + AQ + avPE + nHypSwitch + Variability:Volatility + Variability:TimeBin + Volatility:TimeBin + Variability:AQ + Volatility:AQ + TimeBin:AQ + Variability:Volatility:TimeBin + Variability:Volatility:AQ + Variability:TimeBin:AQ + Volatility:TimeBin:AQ + Variability:Volatility:TimeBin:AQ+( 1 \| id ) |
| AIC | 2893.939 |
| R-squared Marginal | 0.920 |
| R-squared Conditional | 0.964 |

#### Model Results

Fixed Effect Omnibus tests

| | F | Num df | Den df | p |
|---|---|---|---|---|
| Variability | 125.092 | 1 | 511.660 | < .001 |
| Volatility | 6.140 | 1 | 719.046 | 0.013 |
| TimeBin | 252.294 | 4 | 718.606 | < .001 |
| AQ | 0.234 | 1 | 36.500 | 0.632 |
| avPE | 1139.785 | 1 | 486.130 | < .001 |
| nHypSwitch | 8.626 | 1 | 271.145 | 0.004 |
| Variability ✻ Volatility | 10.758 | 1 | 728.613 | 0.001 |
| Variability ✻ TimeBin | 17.938 | 4 | 718.606 | < .001 |
| Volatility ✻ TimeBin | 0.060 | 4 | 718.606 | 0.993 |
| Variability ✻ AQ | 3.443 | 1 | 734.027 | 0.064 |
| Volatility ✻ AQ | 7.034e-6 | 1 | 718.910 | 0.998 |
| TimeBin ✻ AQ | 12.162 | 4 | 718.606 | < .001 |
| Variability ✻ Volatility ✻ TimeBin | 0.445 | 4 | 718.606 | 0.776 |

Fixed Effect Omnibus tests

| | F | Num df | Den df | p |
|---|---|---|---|---|
| Variability ✻ Volatility ✻ AQ | 0.005 | 1 | 718.957 | 0.943 |
| Variability ✻ TimeBin ✻ AQ | 0.138 | 4 | 718.606 | 0.968 |
| Volatility ✻ TimeBin ✻ AQ | 0.032 | 4 | 718.606 | 0.998 |
| Variability ✻ Volatility ✻ TimeBin ✻ AQ | 0.426 | 4 | 718.606 | 0.790 |

Fixed Effects Parameter Estimates

| | | | | 95% Confidence Interval | | | | |
|---|---|---|---|---|---|---|---|---|
| Names | Effect | Estimate | SE | Lower | Upper | df | t | p |
| (Intercept) | (Intercept) | 12.384 | 0.232 | 11.930 | 12.838 | 35.472 | 53.485 | < .001 |
| Variability1 | 2 - 1 | -2.386 | 0.213 | -2.804 | -1.968 | 511.660 | -11.184 | < .001 |
| Volatility1 | 2 - 1 | 0.226 | 0.091 | 0.047 | 0.405 | 719.046 | 2.478 | 0.013 |
| TimeBin1 | 2 - 1 | 0.747 | 0.144 | 0.465 | 1.029 | 718.606 | 5.184 | < .001 |
| TimeBin2 | 3 - 1 | 3.655 | 0.144 | 3.372 | 3.937 | 718.606 | 25.360 | < .001 |
| TimeBin3 | 4 - 1 | -0.006 | 0.144 | -0.289 | 0.276 | 718.606 | -0.043 | 0.966 |
| TimeBin4 | 5 - 1 | -0.221 | 0.144 | -0.504 | 0.061 | 718.606 | -1.537 | 0.125 |
| AQ | AQ | -0.019 | 0.040 | -0.098 | 0.059 | 36.500 | -0.484 | 0.632 |
| avPE | avPE | 1.318 | 0.039 | 1.241 | 1.395 | 486.130 | 33.761 | < .001 |
| nHypSwitch | nHypSwitch | -0.049 | 0.017 | -0.081 | -0.016 | 271.145 | -2.937 | 0.004 |
| Variability1 ✻ Volatility1 | 2 - 1 ✻ 2 - 1 | -0.603 | 0.184 | -0.963 | -0.243 | 728.613 | -3.280 | 0.001 |
| Variability1 ✻ TimeBin1 | 2 - 1 ✻ 2 - 1 | -0.467 | 0.288 | -1.032 | 0.098 | 718.606 | -1.620 | 0.106 |
| Variability1 ✻ TimeBin2 | 2 - 1 ✻ 3 - 1 | -2.158 | 0.288 | -2.723 | -1.593 | 718.606 | -7.488 | < .001 |
| Variability1 ✻ TimeBin3 | 2 - 1 ✻ 4 - 1 | -0.433 | 0.288 | -0.997 | 0.132 | 718.606 | -1.501 | 0.134 |

Fixed Effects Parameter Estimates

| | | | | 95% Confidence Interval | | | | |
|---|---|---|---|---|---|---|---|---|
| **Names** | **Effect** | **Estimate** | **SE** | **Lower** | **Upper** | **df** | **t** | **p** |
| Variability1 ✻ TimeBin4 | 2 - 1 ✻ 5 - 1 | -0.200 | 0.288 | -0.765 | 0.364 | 718.606 | -0.695 | 0.487 |
| Volatility1 ✻ TimeBin1 | 2 - 1 ✻ 2 - 1 | 0.044 | 0.288 | -0.521 | 0.609 | 718.606 | 0.153 | 0.879 |
| Volatility1 ✻ TimeBin2 | 2 - 1 ✻ 3 - 1 | 0.133 | 0.288 | -0.432 | 0.697 | 718.606 | 0.460 | 0.646 |
| Volatility1 ✻ TimeBin3 | 2 - 1 ✻ 4 - 1 | 0.052 | 0.288 | -0.513 | 0.617 | 718.606 | 0.181 | 0.857 |
| Volatility1 ✻ TimeBin4 | 2 - 1 ✻ 5 - 1 | 0.024 | 0.288 | -0.541 | 0.589 | 718.606 | 0.083 | 0.934 |
| Variability1 ✻ AQ | 2 - 1 ✻ AQ | 0.030 | 0.016 | -0.002 | 0.062 | 734.027 | 1.855 | 0.064 |
| Volatility1 ✻ AQ | 2 - 1 ✻ AQ | -4.157e−5 | 0.016 | -0.031 | 0.031 | 718.910 | -0.003 | 0.998 |
| TimeBin1 ✻ AQ | 2 - 1 ✻ AQ | -0.031 | 0.025 | -0.080 | 0.017 | 718.606 | -1.268 | 0.205 |
| TimeBin2 ✻ AQ | 3 - 1 ✻ AQ | -0.144 | 0.025 | -0.192 | -0.095 | 718.606 | -5.804 | < .001 |
| TimeBin3 ✻ AQ | 4 - 1 ✻ AQ | -0.009 | 0.025 | -0.058 | 0.039 | 718.606 | -0.381 | 0.703 |
| TimeBin4 ✻ AQ | 5 - 1 ✻ AQ | -2.090e−5 | 0.025 | -0.049 | 0.049 | 718.606 | -8.436e−4 | 0.999 |
| Variability1 ✻ Volatility1 ✻ TimeBin1 | 2 - 1 ✻ 2 - 1 ✻ 2 - 1 | -0.217 | 0.576 | -1.347 | 0.912 | 718.606 | -0.377 | 0.706 |
| Variability1 ✻ Volatility1 ✻ TimeBin2 | 2 - 1 ✻ 2 - 1 ✻ 3 - 1 | -0.707 | 0.576 | -1.837 | 0.422 | 718.606 | -1.227 | 0.220 |
| Variability1 ✻ Volatility1 ✻ TimeBin3 | 2 - 1 ✻ 2 - 1 ✻ 4 - 1 | -0.420 | 0.576 | -1.549 | 0.710 | 718.606 | -0.728 | 0.467 |

Fixed Effects Parameter Estimates

| | | | | 95% Confidence Interval | | | | |
|---|---|---|---|---|---|---|---|---|
| **Names** | **Effect** | **Estimate** | **SE** | **Lower** | **Upper** | **df** | **t** | **p** |
| Variability1 ✻ Volatility1 ✻ TimeBin4 | 2 - 1 ✻ 2 - 1 ✻ 5 - 1 | -0.164 | 0.576 | -1.294 | 0.966 | 718.606 | -0.284 | 0.776 |
| Variability1 ✻ Volatility1 ✻ AQ | 2 - 1 ✻ 2 - 1 ✻ AQ | 0.002 | 0.031 | -0.059 | 0.064 | 718.957 | 0.071 | 0.943 |
| Variability1 ✻ TimeBin1 ✻ AQ | 2 - 1 ✻ 2 - 1 ✻ AQ | -0.017 | 0.050 | -0.114 | 0.080 | 718.606 | -0.347 | 0.728 |
| Variability1 ✻ TimeBin2 ✻ AQ | 2 - 1 ✻ 3 - 1 ✻ AQ | 0.019 | 0.050 | -0.079 | 0.116 | 718.606 | 0.374 | 0.708 |
| Variability1 ✻ TimeBin3 ✻ AQ | 2 - 1 ✻ 4 - 1 ✻ AQ | 0.002 | 0.050 | -0.095 | 0.099 | 718.606 | 0.033 | 0.974 |
| Variability1 ✻ TimeBin4 ✻ AQ | 2 - 1 ✻ 5 - 1 ✻ AQ | -0.006 | 0.050 | -0.103 | 0.091 | 718.606 | -0.120 | 0.905 |
| Volatility1 ✻ TimeBin1 ✻ AQ | 2 - 1 ✻ 2 - 1 ✻ AQ | -0.007 | 0.050 | -0.104 | 0.090 | 718.606 | -0.147 | 0.883 |
| Volatility1 ✻ TimeBin2 ✻ AQ | 2 - 1 ✻ 3 - 1 ✻ AQ | -0.011 | 0.050 | -0.109 | 0.086 | 718.606 | -0.232 | 0.817 |
| Volatility1 ✻ TimeBin3 ✻ AQ | 2 - 1 ✻ 4 - 1 ✻ AQ | -0.017 | 0.050 | -0.114 | 0.080 | 718.606 | -0.343 | 0.732 |
| Volatility1 ✻ TimeBin4 ✻ AQ | 2 - 1 ✻ 5 - 1 ✻ AQ | -0.012 | 0.050 | -0.109 | 0.086 | 718.606 | -0.233 | 0.816 |
| Variability1 ✻ Volatility1 ✻ TimeBin1 ✻ AQ | 2 - 1 ✻ 2 - 1 ✻ 2 - 1 ✻ AQ | 0.064 | 0.099 | -0.130 | 0.258 | 718.606 | 0.643 | 0.520 |

Fixed Effects Parameter Estimates

| | | | | 95% Confidence Interval | | | | |
|---|---|---|---|---|---|---|---|---|
| **Names** | **Effect** | **Estimate** | **SE** | **Lower** | **Upper** | **df** | **t** | **p** |
| Variability1 ✻ Volatility1 ✻ TimeBin2 ✻ AQ | 2 - 1 ✻ 2 - 1 ✻ 3 - 1 ✻ AQ | 0.126 | 0.099 | -0.068 | 0.320 | 718.606 | 1.271 | 0.204 |
| Variability1 ✻ Volatility1 ✻ TimeBin3 ✻ AQ | 2 - 1 ✻ 2 - 1 ✻ 4 - 1 ✻ AQ | 0.071 | 0.099 | -0.123 | 0.265 | 718.606 | 0.716 | 0.474 |
| Variability1 ✻ Volatility1 ✻ TimeBin4 ✻ AQ | 2 - 1 ✻ 2 - 1 ✻ 5 - 1 ✻ AQ | 0.043 | 0.099 | -0.151 | 0.237 | 718.606 | 0.436 | 0.663 |

Random Components

| Groups | Name | SD | Variance | ICC |
|---|---|---|---|---|
| id | (Intercept) | 1.436 | 2.062 | 0.554 |
| Residual | | 1.289 | 1.661 | |

#### Post Hoc Tests

Post Hoc Comparisons - Variability

| Comparison | | | | | | | |
|---|---|---|---|---|---|---|---|
| **Variability** | | **Variability** | **Difference** | **SE** | **t** | **df** | $p_{bonferroni}$ |
| 1 | - | 2 | 2.386 | 0.215 | 11.094 | 518.881 | < .001 |

Post Hoc Comparisons - Volatility

| Comparison | | | | | | | |
|---|---|---|---|---|---|---|---|
| Volatility | | Volatility | Difference | SE | t | df | $p_{bonferroni}$ |
| 1 | - | 2 | -0.226 | 0.091 | -2.478 | 720.631 | 0.013 |

Post Hoc Comparisons - TimeBin

| Comparison | | | | | | | |
|---|---|---|---|---|---|---|---|
| TimeBin | | TimeBin | Difference | SE | t | df | $p_{bonferroni}$ |
| 2 | - | 3 | -2.908 | 0.144 | -20.176 | 720.208 | < .001 |
| 2 | - | 4 | 0.753 | 0.144 | 5.227 | 720.208 | < .001 |
| 2 | - | 5 | 0.968 | 0.144 | 6.721 | 720.208 | < .001 |
| 1 | - | 2 | -0.747 | 0.144 | -5.184 | 720.208 | < .001 |
| 1 | - | 3 | -3.655 | 0.144 | -25.360 | 720.208 | < .001 |
| 1 | - | 4 | 0.006 | 0.144 | 0.043 | 720.208 | 1.000 |
| 1 | - | 5 | 0.221 | 0.144 | 1.537 | 720.208 | 1.000 |
| 3 | - | 4 | 3.661 | 0.144 | 25.403 | 720.208 | < .001 |
| 3 | - | 5 | 3.876 | 0.144 | 26.897 | 720.208 | < .001 |
| 4 | - | 5 | 0.215 | 0.144 | 1.494 | 720.208 | 1.000 |

Post Hoc Comparisons - Variability ✻ Volatility

| Comparison | | | | | | | | | |
|---|---|---|---|---|---|---|---|---|---|
| Variability | Volatility | | Variability | Volatility | Difference | SE | t | df | $p_{bonferroni}$ |
| 2 | 1 | - | 2 | 2 | 0.076 | 0.129 | 0.584 | 723.906 | 1.000 |
| 2 | 1 | - | 1 | 2 | -2.612 | 0.233 | -11.218 | 584.968 | < .001 |
| 1 | 2 | - | 2 | 2 | 2.687 | 0.239 | 11.223 | 565.458 | < .001 |

Post Hoc Comparisons - Variability ✻ Volatility

| Comparison | | | | | | | | | |
|---|---|---|---|---|---|---|---|---|---|
| Variability | Volatility | | Variability | Volatility | Difference | SE | t | df | $p_{bonferroni}$ |
| 1 | 1 | - | 2 | 2 | 2.160 | 0.234 | 9.217 | 589.917 | < .001 |
| 1 | 1 | - | 2 | 1 | 2.084 | 0.228 | 9.135 | 608.443 | < .001 |
| 1 | 1 | - | 1 | 2 | -0.527 | 0.130 | -4.068 | 726.869 | < .001 |

Post Hoc Comparisons - Variability ✻ TimeBin

| Comparison | | | | | | | | | |
|---|---|---|---|---|---|---|---|---|---|
| Variability | TimeBin | | Variability | TimeBin | Difference | SE | t | df | $p_{bonferroni}$ |
| 2 | 2 | - | 2 | 3 | -2.062 | 0.204 | -10.117 | 720.208 | < .001 |
| 2 | 2 | - | 2 | 4 | 0.736 | 0.204 | 3.612 | 720.208 | 0.015 |
| 2 | 2 | - | 2 | 5 | 0.835 | 0.204 | 4.099 | 720.208 | 0.002 |
| 2 | 2 | - | 1 | 3 | -5.954 | 0.282 | -21.121 | 697.056 | < .001 |
| 2 | 2 | - | 1 | 4 | -1.431 | 0.282 | -5.075 | 697.056 | < .001 |
| 2 | 2 | - | 1 | 5 | -1.099 | 0.282 | -3.900 | 697.056 | 0.005 |
| 2 | 1 | - | 2 | 2 | -0.514 | 0.204 | -2.520 | 720.208 | 0.537 |
| 2 | 1 | - | 2 | 3 | -2.575 | 0.204 | -12.638 | 720.208 | < .001 |
| 2 | 1 | - | 2 | 4 | 0.222 | 0.204 | 1.091 | 720.208 | 1.000 |
| 2 | 1 | - | 2 | 5 | 0.322 | 0.204 | 1.578 | 720.208 | 1.000 |
| 2 | 1 | - | 1 | 2 | -2.715 | 0.282 | -9.630 | 697.056 | < .001 |
| 2 | 1 | - | 1 | 3 | -6.468 | 0.282 | -22.943 | 697.056 | < .001 |
| 2 | 1 | - | 1 | 4 | -1.944 | 0.282 | -6.897 | 697.056 | < .001 |
| 2 | 1 | - | 1 | 5 | -1.613 | 0.282 | -5.722 | 697.056 | < .001 |
| 2 | 3 | - | 2 | 4 | 2.798 | 0.204 | 13.729 | 720.208 | < .001 |
| 2 | 3 | - | 2 | 5 | 2.897 | 0.204 | 14.216 | 720.208 | < .001 |

Post Hoc Comparisons - Variability ✻ TimeBin

| Comparison | | | | | | | | | |
|---|---|---|---|---|---|---|---|---|---|
| Variability | TimeBin | | Variability | TimeBin | Difference | SE | t | df | $p_{bonferroni}$ |
| 2 | 3 | - | 1 | 4 | 0.631 | 0.282 | 2.239 | 697.056 | 1.000 |
| 2 | 3 | - | 1 | 5 | 0.962 | 0.282 | 3.414 | 697.056 | 0.030 |
| 2 | 4 | - | 2 | 5 | 0.099 | 0.204 | 0.487 | 720.208 | 1.000 |
| 2 | 4 | - | 1 | 5 | -1.835 | 0.282 | -6.511 | 697.056 | < .001 |
| 1 | 2 | - | 2 | 2 | 2.201 | 0.282 | 7.808 | 697.056 | < .001 |
| 1 | 2 | - | 2 | 3 | 0.139 | 0.282 | 0.494 | 697.056 | 1.000 |
| 1 | 2 | - | 2 | 4 | 2.937 | 0.282 | 10.419 | 697.056 | < .001 |
| 1 | 2 | - | 2 | 5 | 3.036 | 0.282 | 10.771 | 697.056 | < .001 |
| 1 | 2 | - | 1 | 3 | -3.753 | 0.204 | -18.416 | 720.208 | < .001 |
| 1 | 2 | - | 1 | 4 | 0.770 | 0.204 | 3.780 | 720.208 | 0.008 |
| 1 | 2 | - | 1 | 5 | 1.102 | 0.204 | 5.406 | 720.208 | < .001 |
| 1 | 1 | - | 2 | 2 | 1.221 | 0.282 | 4.330 | 697.056 | < .001 |
| 1 | 1 | - | 2 | 1 | 1.734 | 0.282 | 6.152 | 697.056 | < .001 |
| 1 | 1 | - | 2 | 3 | -0.841 | 0.282 | -2.984 | 697.056 | 0.132 |
| 1 | 1 | - | 2 | 4 | 1.957 | 0.282 | 6.941 | 697.056 | < .001 |
| 1 | 1 | - | 2 | 5 | 2.056 | 0.282 | 7.293 | 697.056 | < .001 |
| 1 | 1 | - | 1 | 2 | -0.980 | 0.204 | -4.811 | 720.208 | < .001 |
| 1 | 1 | - | 1 | 3 | -4.734 | 0.204 | -23.227 | 720.208 | < .001 |
| 1 | 1 | - | 1 | 4 | -0.210 | 0.204 | -1.031 | 720.208 | 1.000 |
| 1 | 1 | - | 1 | 5 | 0.121 | 0.204 | 0.595 | 720.208 | 1.000 |
| 1 | 3 | - | 2 | 3 | 3.892 | 0.282 | 13.807 | 697.056 | < .001 |
| 1 | 3 | - | 2 | 4 | 6.690 | 0.282 | 23.732 | 697.056 | < .001 |
| 1 | 3 | - | 2 | 5 | 6.789 | 0.282 | 24.084 | 697.056 | < .001 |
| 1 | 3 | - | 1 | 4 | 4.523 | 0.204 | 22.196 | 720.208 | < .001 |
| 1 | 3 | - | 1 | 5 | 4.855 | 0.204 | 23.822 | 720.208 | < .001 |

Post Hoc Comparisons - Variability ✻ TimeBin

| Comparison | | | | | | | | | |
|---|---|---|---|---|---|---|---|---|---|
| Variability | TimeBin | | Variability | TimeBin | Difference | SE | t | df | $p_{bonferroni}$ |
| 1 | 4 | - | 2 | 4 | 2.167 | 0.282 | 7.686 | 697.056 | < .001 |
| 1 | 4 | - | 2 | 5 | 2.266 | 0.282 | 8.038 | 697.056 | < .001 |
| 1 | 4 | - | 1 | 5 | 0.331 | 0.204 | 1.626 | 720.208 | 1.000 |
| 1 | 5 | - | 2 | 5 | 1.935 | 0.282 | 6.863 | 697.056 | < .001 |

#### Simple Effects

Simple effects of AQ : Omnibus Tests

| Moderator levels | | | | |
|---|---|---|---|---|
| TimeBin | F | Num df | Den df | p |
| 1 | 0.165 | 1.000 | 50.450 | 0.686 |
| 2 | 0.104 | 1.000 | 50.450 | 0.748 |
| 3 | 8.576 | 1.000 | 50.450 | 0.005 |
| 4 | 0.035 | 1.000 | 50.450 | 0.852 |
| 5 | 0.165 | 1.000 | 50.450 | 0.687 |

Simple effects of AQ : Parameter estimates

| Moderator levels | | | 95% Confidence Interval | | | | |
|---|---|---|---|---|---|---|---|
| TimeBin | Estimate | SE | Lower | Upper | df | t | p |
| 1 | 0.018 | 0.043 | -0.069 | 0.104 | 50.450 | 0.406 | 0.686 |
| 2 | -0.014 | 0.043 | -0.100 | 0.073 | 50.450 | -0.322 | 0.748 |
| 3 | -0.126 | 0.043 | -0.213 | -0.040 | 50.450 | -2.928 | 0.005 |
| 4 | 0.008 | 0.043 | -0.079 | 0.095 | 50.450 | 0.187 | 0.852 |

Simple effects of AQ : Parameter estimates

| Moderator levels | | | 95% Confidence Interval | | | | |
|---|---|---|---|---|---|---|---|
| TimeBin | Estimate | SE | Lower | Upper | df | t | p |
| 5 | 0.017 | 0.043 | -0.069 | 0.104 | 50.450 | 0.406 | 0.687 |

Note. Simple effects are estimated setting higher order moderator (if any) in covariates to zero and averaging across moderating factors levels (if any)

#### Correlation Matrix

Correlation Matrix

| | | AQ | HypSwitchERPE |
|---|---|---|---|
| AQ | Pearson's r | — | |
| | p-value | — | |
| HypSwitchERPE | Pearson's r | -0.211 | — |
| | p-value | < .001 | — |

#### Plot

AQ HypSwitchERPE

AQ

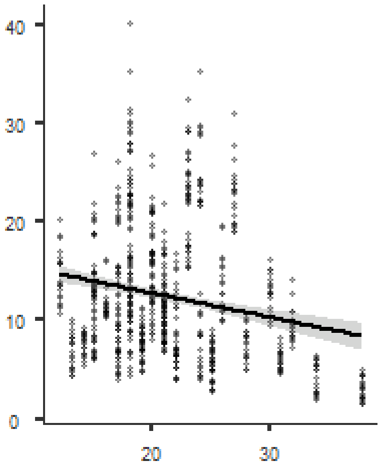

HypSwitchERPE
